# Functional connectivity favors aberrant visual network c-Fos expression accompanied by cortical synapse loss in a mouse model of Alzheimer’s disease

**DOI:** 10.1101/2023.01.05.522900

**Authors:** Oliver J. L’Esperance, Josh McGhee, Garett Davidson, Suraj Niraula, Adam Smith, Alexandre A Sosunov, Shirley Shidu Yan, Jaichandar Subramanian

## Abstract

While Alzheimer’s disease (AD) has been extensively studied with a focus on cognitive networks, sensory network dysfunction has received comparatively less attention despite compelling evidence of its significance in both Alzheimer’s disease patients and mouse models. We recently found that neurons in the primary visual cortex of an AD mouse model expressing human amyloid protein precursor with the Swedish and Indiana mutations (hAPP mutations) exhibit aberrant c-Fos expression and altered synaptic structures at a pre-amyloid plaque stage. However, it is unclear whether aberrant c-Fos expression and synaptic pathology vary across the broader visual network and to what extent c-Fos abnormality in the cortex is inherited through functional connectivity. Using both sexes of 4-6-month AD model mice with hAPP mutations (J20[PDGF-APPSw, Ind]), we found that cortical regions of the visual network show aberrant c-Fos expression and impaired experience-dependent modulation while subcortical regions do not. Interestingly, the average network-wide functional connectivity strength of a brain region in wild type (WT) mice significantly predicts its aberrant c-Fos expression, which in turn correlates with impaired experience-dependent modulation in the AD model. Using *in vivo* two-photon and *ex vivo* imaging of presynaptic termini, we observed a subtle yet selective weakening of excitatory cortical synapses in the visual cortex. Intriguingly, the change in the size distribution of cortical boutons in the AD model is downscaled relative to those in WT mice, suggesting that synaptic weakening may reflect an adaptation to aberrant activity. Our observations suggest that cellular and synaptic abnormalities in the AD model represent a maladaptive transformation of the baseline physiological state seen in WT conditions rather than entirely novel and unrelated manifestations.

## Introduction

Accumulating evidence shows cellular and synaptic loss within the visual cortex in individuals affected by Alzheimer’s disease (AD) (Armstrong, 2009; Beach et al., 1989; Cui et al., 2007; Frisoni et al., 2009; Hill et al., 2014; Hof and Morrison, 1990; Hwang et al., 2021; Kurucu et al., 2022; Leuba and Kraftsik, 1994; Lewis et al., 1987; Mavroudis et al., 2011; Metsaars et al., 2003; Morrison et al., 1991). AD patients exhibit visual impairments in contrast sensitivity, acuity, depth perception, and facial recognition (Cormack et al., 2000; Kirby et al., 2010; Kusne et al., 2017; Mandal et al., 2012; Murphy, 2019; Zhang et al., 2023). In some patients, visual system disruption and visuospatial deficits emerge early in the disease progression, particularly in those developing the posterior cortical atrophy variant of AD (Best et al., 2023; Chen et al., 2019; Crutch et al., 2012; Holden et al., 2020; Maia da Silva et al., 2017; North et al., 2021; Yerstein et al., 2021). In these patients, cortical areas of the visual system frequently display irregular patterns of activation and functional connectivity (Fredericks et al., 2019; Migliaccio et al., 2016). Mouse models of AD have also shown structural and functional visual system disruption (Chen et al., 2022; Chiquita et al., 2019; Criscuolo et al., 2018; Grienberger et al., 2012; Liebscher et al., 2016; Maya-Vetencourt et al., 2014; Niraula et al., 2023a; Niraula et al., 2024; Rosales Jubal et al., 2021; Rudinskiy et al., 2012; Vit et al., 2021; William et al., 2012; William et al., 2021). Visual impairments contribute to cognitive deficits in AD patients (Lad et al., 2024; Rizzo et al., 2000), and disruption to visual cortical regions impairs performance in cognitive tasks, such as the Morris water maze (Conejo et al., 2007; Espinoza et al., 1999; Hoh et al., 2003; Prusky et al., 2000) in rodents.

We recently observed an increase in the expression of the immediate early gene c-Fos in the primary visual cortex of an AD mouse model (Niraula et al., 2023a). c-Fos is rapidly and transiently expressed following membrane depolarization and calcium influx in some neurons (Kawashima et al., 2014; Mahringer et al., 2022; Morgan et al., 1987; Zangenehpour and Chaudhuri, 2002). Its low baseline expression, widespread expression profile, and close associations with learning and memory recall have made c-Fos^+^ cell labeling an attractive tool for mapping and manipulating experience and memory-associated neuronal ensembles (Barth et al., 2004; Guenthner et al., 2013; Liu et al., 2012; Reijmers et al., 2007). c-Fos^+^ neurons fire more frequently and are more likely to share direct and indirect connections than c-Fos^-^ neurons (Yassin et al., 2010). Due to these properties, correlated c-Fos expression between brain regions has frequently been used to represent functionally connected networks in a variety of contexts (Borcuk et al., 2022; Silva et al., 2019; Tanimizu et al., 2017; Teles et al., 2015; Vetere et al., 2017; Wheeler et al., 2013). Our recent observations of elevated visual cortical c-Fos expression, altered postsynaptic structures, and impaired visual experience-dependent plasticity and visual memory in a pre-amyloid plaque stage AD mouse model are suggestive of a structurally and functionally impaired visual network (Niraula et al., 2023a; Niraula et al., 2024). However, it is unclear whether disruption to the visual network at this early stage is localized to specific regions within the network and whether functional connectivity between these regions plays a role. Addressing these questions would shed light on early sensory network abnormalities that could impair visuospatial perception and cognition without overt plaque accumulation or neurodegeneration. Here, we addressed these questions using pre-amyloid plaque (5-6-months-old) stage mice bearing Swedish and Indiana mutations in the human amyloid precursor protein (J20 line (hAPP mice); (Mucke et al., 2000) with human-like early-stage cortical amyloid deposition (Whitesell et al., 2019).

## Results

### Aberrant c-Fos+ cell distribution emerges in cortical regions of the visual network in hAPP mice

To test whether increased c-Fos^+^ cell density in the primary visual cortex of 4-6-month-old J20-hAPP mice (Niraula et al., 2023a) is a feature of the entire visual network or the brain at large, we performed c-Fos immunohistochemistry on 24 evenly spaced coronal sections spanning the brain of hAPP mice and WT littermate controls and mapped them to the Allen Mouse Brain CCFv3 for regional analysis (Figure 1A-D, see Supplementary Table 1 for region abbreviations). Cells were automatically counted with consistent parameters for size range (8-18 µm diameter), circular drop-off in fluorescence as determined by an omnidirectional rolling Laplacian of Gaussian (LoG) transformation of pixel values, and a local brightness threshold (steepness of LoG), which was set such that dim cells with near background c-Fos expression and false positives arising from background fluorescence fluctuation (∼70% of all detected 8-18 µm diameter “objects” in WT sections) were excluded (Supplementary Figure 1A-B). ∼90% of c-Fos^+^ cells identified with these parameters colocalized with NeuN^+^ cells with no difference in colocalization between genotypes (Supplementary Figure 1C-D).

**Figure 1:**
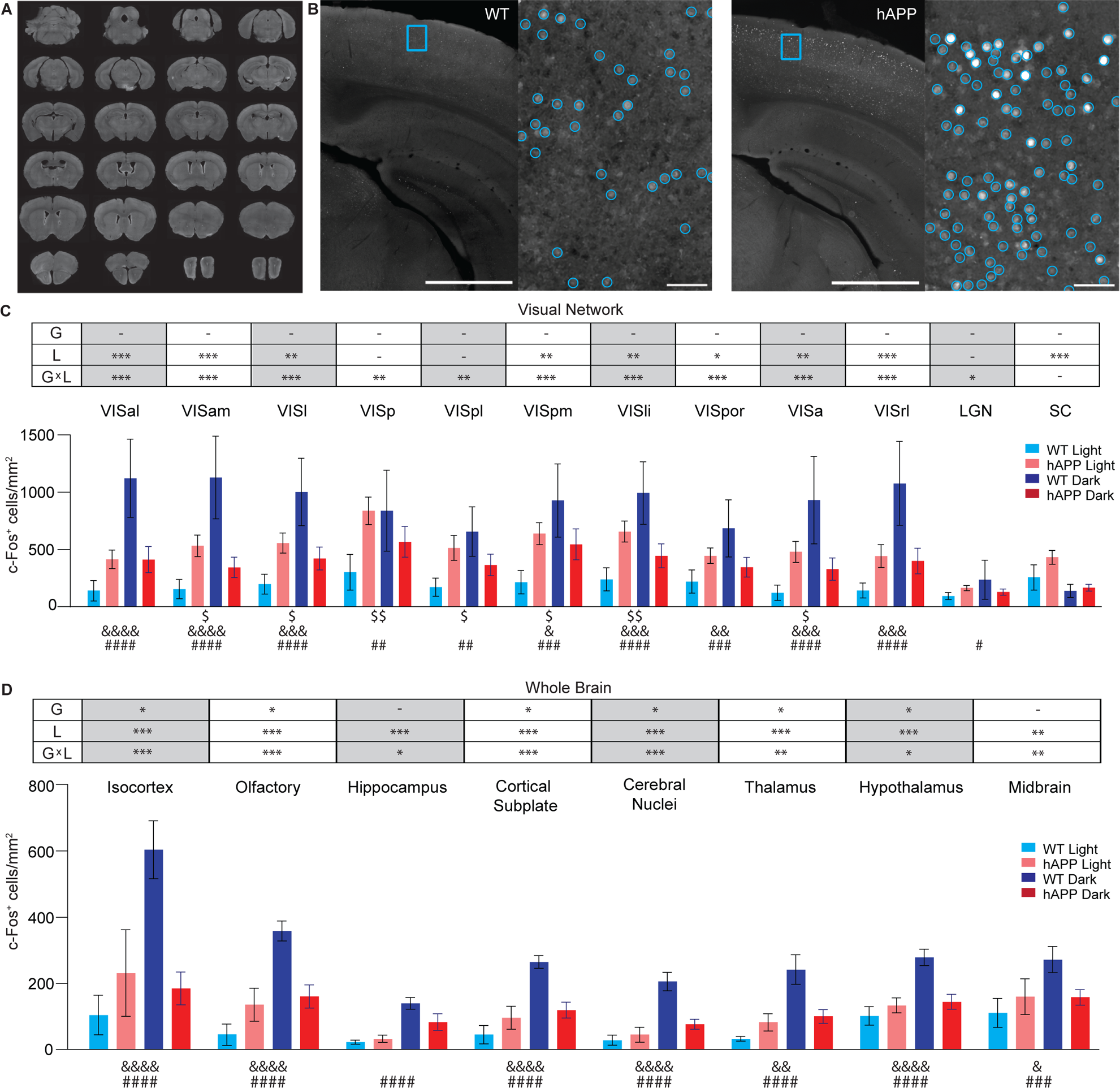
hAPP mice exhibit high baseline c-Fos^+^ cell density and reduced c-Fos modulation following light deprivation. **A)** Representative images of 24 evenly spaced landmark sections used for brain-wide c-Fos^+^ cell density quantification. **B)** Representative images of c-Fos fluorescence in the primary visual cortex of wildtype (WT, left) and hAPP mice (right). The area in the blue box is shown in higher magnification in the right half of each panel. Blue circles indicate c-Fos^+^ cells identified by automatic cell detection. The identification threshold was set such that dim cells representing close to background c-Fos expression and false positives from background fluorescence fluctuation (∼70% of all detected “objects” in WT sections) were excluded. Scale bar: 1mm (left half of each panel), 100µm (right half). **C-D)** c-Fos^+^ cell density across regions of the visual network (C) areas across the brain (D) for WT and hAPP mice under a standard 12 h light/dark cycle (Light) and one week of light deprivation (Dark). 2-way ANOVAs (genotype, light experience, interaction), with Benjamini-Hochberg’s false discovery rate correction for brain regions shown above. Post-hoc comparisons (Šídák’s multiple comparisons test, 1 family with 4 comparisons) are shown for regions with a significant interaction. Benjamini-Hochberg corrected p-values: -p > 0.05, *p < 0.05, **p < 0.01, ***p < 0.001. F(1, 31) for all factors in all ANOVAs. Corrected p-value for post-hoc comparisons: WT Light vs. hAPP Light: ^$^p < 0.05, ^$$^p < 0.01, WT Dark vs. hAPP Dark: ^&^p < 0.05, ^&&^p < 0.01, ^&&&^p < 0.001, ^&&&&^p < 0.0001, WT Light vs. WT Dark: ^#^p < 0.05, ^##^p < 0.01, ^###^p < 0.001, ^####^p < 0.0001. Perfusions for both light and dark groups were performed close to the center of the light phase in the standard 12 h light/dark cycle to avoid circadian transition-related effects. Data are presented as mean ± SEM. n = 8 WT-light, 9 WT-dark, 10 hAPP-light, and 8 hAPP-dark mice.

We confirmed that the primary visual cortex exhibited increased c-Fos^+^ cell density in hAPP mice housed under a 12-hour light/12-hour dark cycle compared to WT mice housed under the same conditions (hAPP-light and WT-light mice, respectively; Figure 1C). However, the increase was not limited to the primary visual cortex, with multiple higher visual areas exhibiting increased c-Fos^+^ cell density (Figure 1C). Strikingly, the primary visual cortex displayed the densest c-Fos^+^ cells of any region in WT mice (301.4 ± 54.6 cells/mm^2^) and the largest average magnitude of increased c-Fos^+^ cell density in hAPP mice (+536.8 cells/mm^2^, Figure 1C). Within the primary visual cortex, we found that all layers containing neuronal cell bodies show elevated c-Fos^+^ cell density in hAPP-light compared to WT-light mice (Supplementary Figure 2A). In contrast, the lateral geniculate nucleus, a subcortical visual region upstream of the primary visual cortex, and the superior colliculus did not exhibit significant differences between the genotypes Figure 1C).

To investigate the contribution of light experience to aberrant c-Fos expression in the hAPP visual network, we housed WT and hAPP mice in complete darkness (24-hour dark cycle) for one week (WT-dark and hAPP-dark, respectively) and quantified the resulting c-Fos^+^ cell density (Figure 1C-D, Supplementary Figure 2, Table 1). Interestingly, c-Fos^+^ cell density was significantly increased in WT-dark mice across cortical areas of the visual system (Figure 1C, Supplementary Figure 2A). In contrast, c-Fos^+^ density was not significantly changed in hAPP-dark mice, with the exception of reductions in c-Fos^+^ cell density in the superior colliculus (Figure 1C). Overall, we observed significant interactions between genotype (G), light experience (L), and brain region (B) across the visual network (B×G×L: F[11, 340] p < 0.0001, 3-way mixed model restricted maximum likelihood ANOVA). Individual brain regions in the visual network showed a significant effect of light experience and its interaction with the genotype (G: F[1, 31] p = 0.44, L: F[1, 31] p = 0.003, G×L: F[1, 31] p < 0.0001, 2-way ANOVA with Benjamini-Hochberg false discovery rate correction for multiple comparisons).

**Table 1:**
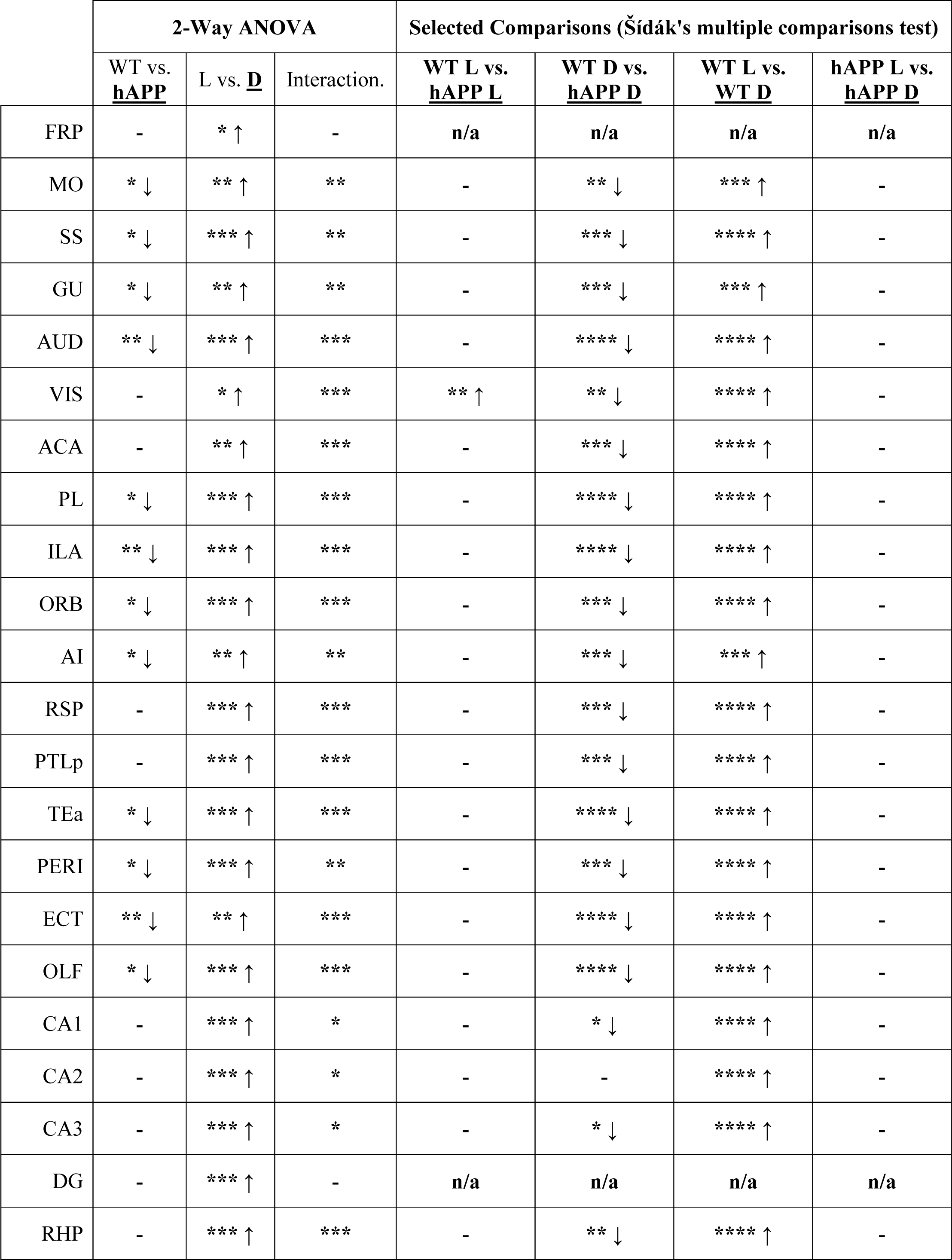
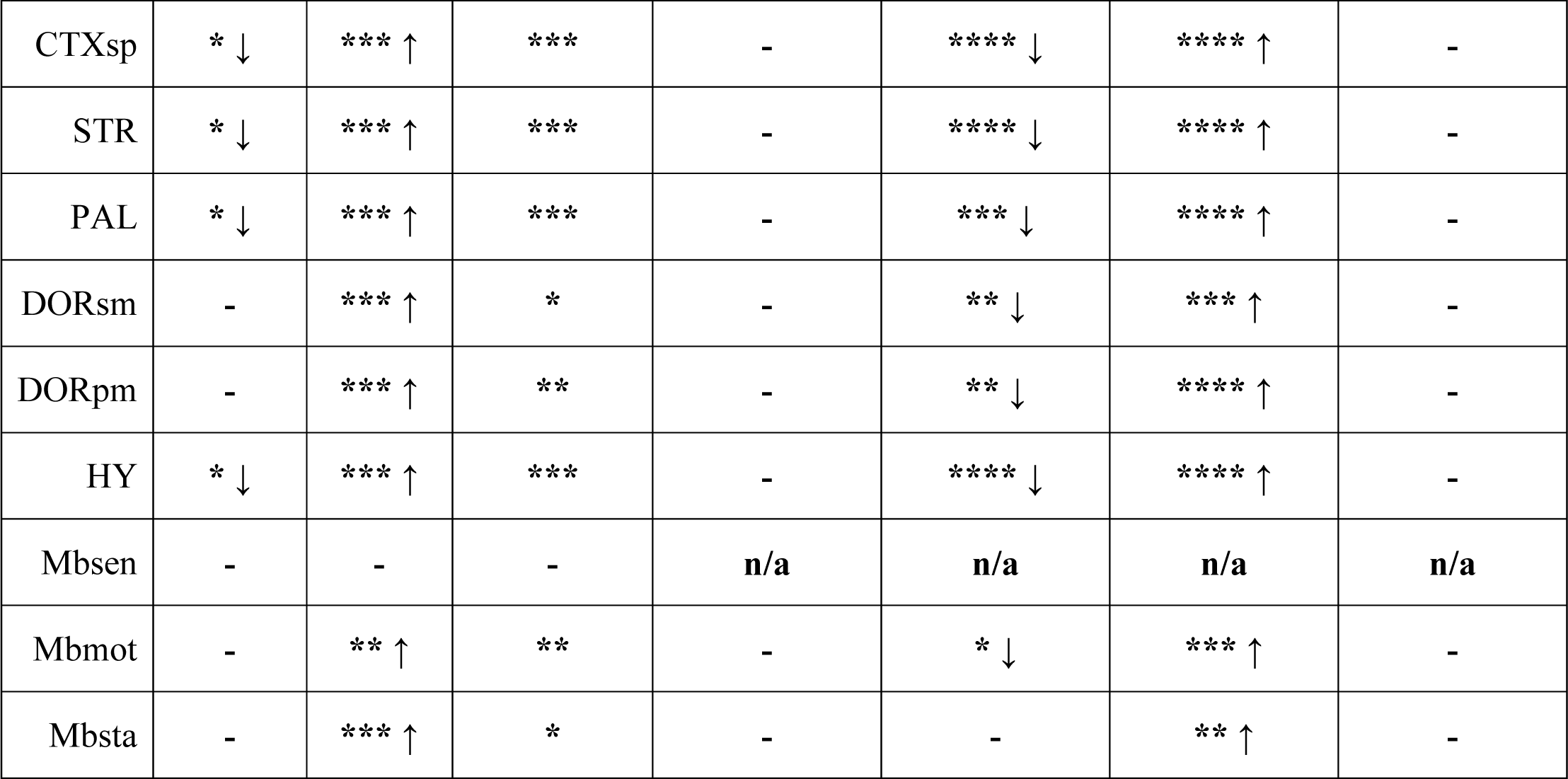
Whole brain mixed-effects ANOVA of c-Fos^+^ cell density with selected comparisons. . 2-way ANOVAs (WT/hAPP [between], Light/Dark [between], interaction [between]), with Benjamini-Hochberg’s false discovery rate correction for brain regions shown. Selected post-hoc comparisons (Šídák’s multiple comparisons test, 1 family with 4 comparisons) are shown for regions with a significant interaction. Directionality of mean c-Fos^+^ cell density change is indicated for the underlined factor/group. Perfusions for both Light and Dark groups were performed close to the center of the light phase in the standard 12h light/dark cycle to avoid circadian transition-related effects. n = 8 WT light, 9 WT dark, 10 hAPP light, and 8 hAPP dark housed mice. F(1,31) for all factors of all ANOVAs. Corrected p-values: -p > 0.05, *p < 0.05, **p < 0.01, ***p < 0.001, ****p < 0.0001.

Outside of the visual network, c-Fos^+^ cell density was not significantly different between WT-light and hAPP-light mice after correcting for multiple comparisons in any brain region (Figure 1D), though parietal (3.5x) and retrosplenial (3.1x) cortices, which are adjacent to visual cortices, showed a large increase in hAPP-light mice (Supplementary Figure 2, Table 1). Interestingly, light deprivation significantly increased c-Fos^+^ cell density outside the visual network in WT-dark but not hAPP-dark mice compared to light-housed mice of the same genotype (Figure 1D, Supplementary Figure 2, Table 1). Overall, as in the visual network, c-Fos^+^ cell density was influenced by the interaction of brain region, genotype, and light condition (B×G L: all F[30, 930] p < 0.0001, 3-way mixed model restricted maximum likelihood ANOVA). Individual brain regions in the whole brain network showed a significant effect of genotype, light experience, and their interaction (G: F[1, 31] p = 0.008, L: F[1, 31] p < 0.0001, G×L: F[1, 31] p < 0.0001, 2-way ANOVA with Benjamini-Hochberg false discovery rate correction for multiple comparisons.

Increased c-Fos^+^ cell density in WT-dark mice may partially result from heightened anxiety due to prolonged light deprivation or circadian shifts. We thus investigated home cage activity patterns and behavioral anxiety responses for WT-light and WT-dark mice (Supplementary Figure 3A). We found that following one week of light deprivation, mice tended to become active slightly earlier than they normally would under 12-hour light/12-hour dark housing conditions (Supplementary Figure 3B) in line with previous reports (Azzi et al., 2014; Eckel-Mahan and Sassone-Corsi, 2015; Mistlberger and Holmes, 2000; Valentinuzzi et al., 1997). However, when these mice were introduced to an open field chamber after one week of light deprivation, they did not spend less time exploring the center of a dark or lit open field chamber, a measure of anxiety-like behavior (Gould et al., 2009; Seibenhener and Wooten, 2015), than their standard light housed controls (Supplementary Figure 3C-D).

We next tested if the c-Fos^+^ cell density increase in the primary visual cortex of WT mice during long-term light deprivation is also present under short-term deprivation to minimize the potential effect of circadian drift (Supplementary Figure 3B). To do so, hAPP and WT mice were dark-housed for a total of 40 continuous hours rather than one full week, then compared primary visual cortex c-Fos^+^ cell density between genotypes. The primary visual cortex of WT mice (318.50 ± 86.71 c-Fos^+^ cells/mm^2^) showed nearly a 2-fold increase over the density observed in the hAPP primary visual cortex (115.60 ± 22.46 c-Fos^+^ cells/mm^2^). Though this trend did not reach statistical significance (p = 0.059, student’s *t*-test with Welch’s correction; n = 7 WT and 5 hAPP mice), the variance was nearly four times as high in the WT group (p = 0.011, F test to compare variances), indicating that short term light deprivation increases c-Fos in some but not all WT mice.

### Functional connectivity favors aberrant c-Fos^+^ cell density across the brain of hAPP mice

Correlated c-Fos^+^ cell density in a pair of regions indicates functional connectivity between those regions (Silva et al., 2019; Tanimizu et al., 2017; Teles et al., 2015; Vetere et al., 2017; Wheeler et al., 2013). Previous studies have shown that hyperconnectivity is associated with excessive neuronal activation in early AD networks (Corriveau-Lecavalier et al., 2021; Koelewijn et al., 2019). To investigate whether hAPP mice display hyperconnectivity, we calculated functional connectivity strength for every pair of imaged regions (Pearson’s r of c-Fos^+^ density) both within the visual network and across the brain. Visual network (Figure 2A, left) functional connectivity matrices revealed an increase in pairwise correlation strength between brain regions in hAPP-light (lower) relative to WT-light (upper) mice. One week of light deprivation, which increases c-Fos^+^ density, also increased functional connectivity across the WT-dark visual network, including subcortical regions (Figure 2B, upper left). In contrast, hAPP-dark visual network functional connectivity was largely unaltered (Figure 2B, lower left). Overall, the average functional connectivity of the visual network, quantified as the mean of z values (Fisher’s r-to-z transformation) of each brain region in the network, showed a significant effect of genotype, light experience, and their interaction (G×L: F[1, 22], p < 0.001, 2-way mixed effects restricted maximum likelihood ANOVA, Figure 2C, left). We extended this analysis across the entire brain and observed a significant effect of genotype and light experience but not their interaction. (G×L: F[1, 60], p > 0.05, 2-way mixed effects restricted maximum likelihood ANOVA, Figure 2C, right). Intriguingly, the increase in dark-associated connectivity in the hAPP whole brain network was qualitatively different from that of the WT network. In the WT network, visual deprivation increased functional connectivity between some lateral cortical and subcortical regions, whereas in hAPP mice, it was distributed across nearly all cortical and subcortical regions (Figure 2B).

**Figure 2:**
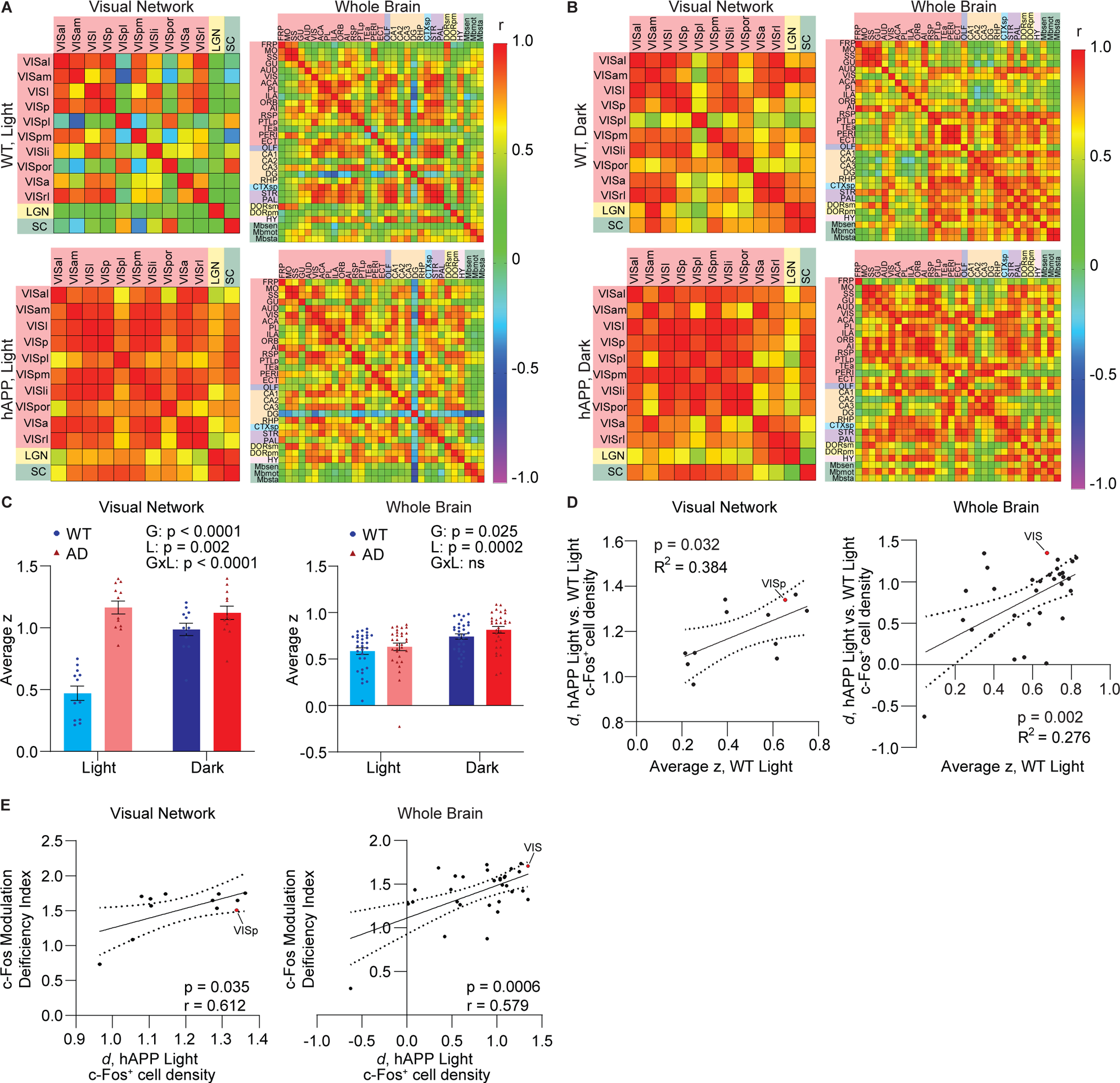
Functional connectivity in WT networks correlates with aberrant c-Fos expression. **A)** Functional connectivity matrices showing pairwise correlation (Pearson’s *r*) of c-Fos^+^ density between regions in the visual (left) and the brain-wide (right) networks for WT (upper) and hAPP (lower) mice reared under 12 h light/12 h dark conditions (Light). Region label color indicates anatomical brain area. Warmer matrix colors indicate stronger pairwise functional connections between regions. **B)** Visual network (left) and brain-wide (right) functional connectivity matrices for WT (upper) and hAPP (lower) mice following one week of light deprivation (Dark). C) Each region’s average correlation strength (average z following r-to-z transformation) across the visual (left) and whole brain networks (right) in WT and hAPP mice for light and dark conditions. Each data point represents the average functional connectivity strength (z) of one brain region across its respective network. 2-way mixed effects restricted maximum likelihood ANOVA with repeated measures (genotype [G], light exposure [L], interaction [G×L], brain region matching). Data are presented as mean ± SEM. ns, not significant (p > 0.05). D) Pearson’s correlation (*r*) of each region’s average z across WT networks (visual, left; whole brain, right) with the effect size (Cohen’s *d*) of hAPP vs. WT (Light groups) on c-Fos^+^ cell density. **E)** Pearson’s correlation (*r*) of each region’s effect size of hAPP (Light) on c-Fos^+^ cell density with its c-Fos modulation deficiency index. Each data point represents one region’s average value. VISp (visual network) and VIS (whole brain network) data points are indicated for D-F. n = 8 WT light, 9 WT dark, 10 hAPP light, and 8 hAPP dark housed mice for all figures. Dotted curves represent 95% confidence intervals. R^2^ represents the coefficient of determination

Abnormal c-Fos^+^ cell density, observed in multiple brain regions of hAPP-light mice (Figure 1C-D, Supplementary Figure 2), may emerge independently or may arise, in part, due to their functional connections with other regions. Thus, we next investigated whether a given region’s average network connection strength in WT-light mice is predictive of the increase in its c-Fos^+^ cell density in hAPP-light mice. We correlated each region’s average connection strength across the WT visual or brain-wide network (average z) with the effect size of hAPP overexpression on c-Fos^+^ density for that region (Cohen’s *d*, see Supplementary Figure 4-5 for all effect sizes). A significant positive correlation emerged between these factors, both within the visual (p = 0.02, Pearson’s correlation, Figure 2D, left) and the whole brain (p = 0.002, Pearson’s correlation, Figure 2D, right) networks. Furthermore, WT-light functional connection strength explained ∼38% and ∼28% of the variance in the effect size of c-Fos^+^ density in the hAPP-light visual network and the whole brain, respectively.

Regions with aberrant c-Fos expression may also show deficiencies in experience-dependent modulation of c-Fos expression in hAPP mice. To test their relationship, we calculated a c-Fos modulation deficiency index (described in Methods). Briefly, the c-Fos modulation deficiency index represents the effect size (absolute value of Cohen’s *d*) of the difference in fold-change of c-Fos expression elicited by light deprivation in WT and hAPP mice. Brain regions with higher c-Fos modulation deficiency index show weaker modulation of c-Fos experience by light deprivation in hAPP mice relative to their WT controls. We found that the effect size of aberrant c-Fos expression in hAPP-light mice positively correlates with c-Fos modulation deficiency index in both the visual (p = 0.035, Pearson’s correlation, Figure 2E, left) and the whole brain (p = 0.0006, Pearson’s correlation, Figure 2E, right) networks, indicating that brain regions with aberrant c-Fos expression show decreased experience-dependent modulation. The reduced experience-dependent modulation in high c-Fos-expressing brain regions of hAPP mice is not due to a general ceiling effect for c-Fos expression because these regions exhibit even higher average c-Fos^+^ density in WT-dark mice in every grouped brain area (67 - 326% increase compared to hAPP-dark mice, Figure 1D).

### hAPP overexpression selectively weakens an excitatory presynaptic terminal subtype in the visual cortex

The cortex-specific increase in the c-Fos expression in the visual network led us to test whether synaptic changes in hAPP mice also exhibit a regional difference. To this end, we immunolabeled excitatory presynaptic vesicular transporters (VGluT1 and VGluT2) and inhibitory presynaptic vesicular transporters (VGAT) in two visual network regions of WT-light mice: the primary visual cortex and the dorsal part of the lateral geniculate nucleus (see Supplementary Figure 6A for the quantification of low magnification images). Though we observed some colocalized VGluT1^+^ and VGluT2^+^ puncta, similar to previous studies (Graziano et al., 2008; Herzog et al., 2006; Li et al., 2003; Nakamura et al., 2008), we analyzed them separately in their respective channels. Semiautomated puncta identification (Figure 3A, Supplementary Figure 6B-C, expected size range 0.32-1.97 µm diameter) revealed a significant decrease in VGluT1^+^ (p < 0.05, student’s *t-*test) but not VGluT2^+^ or VGAT^+^ puncta densities in the primary visual cortex of hAPP-light compared to WT-light mice (Figure 3B-D). In contrast, the dorsal part of the lateral geniculate nucleus showed no significant changes in any puncta densities between the genotypes (Figure 3B-D). Large puncta outside of the expected size range that represented a cluster of 2-3 puncta were occasionally automatically marked as a single punctum. Therefore, we manually counted instances of these undercounted clusters in a 10 µm wide column spanning all layers of the primary visual cortex to estimate the fraction of undercounted clusters. We found that ∼5% or less of puncta in each channel were undercounted due to clustering, and clustering fractions did not differ between genotypes (Figure 3E-G). The lack of significant reductions in VGAT and VGluT2 puncta in hAPP-light compared to WT-light mice was further validated with semiautomated analysis of 3D reconstructions of synaptic puncta labeling obtained using high-resolution confocal microscopy (Supplementary Figure 6D-E). Note that the puncta densities do not reflect the absolute number of synapses but rather indicate relative differences between genotypes.

**Figure 3:**
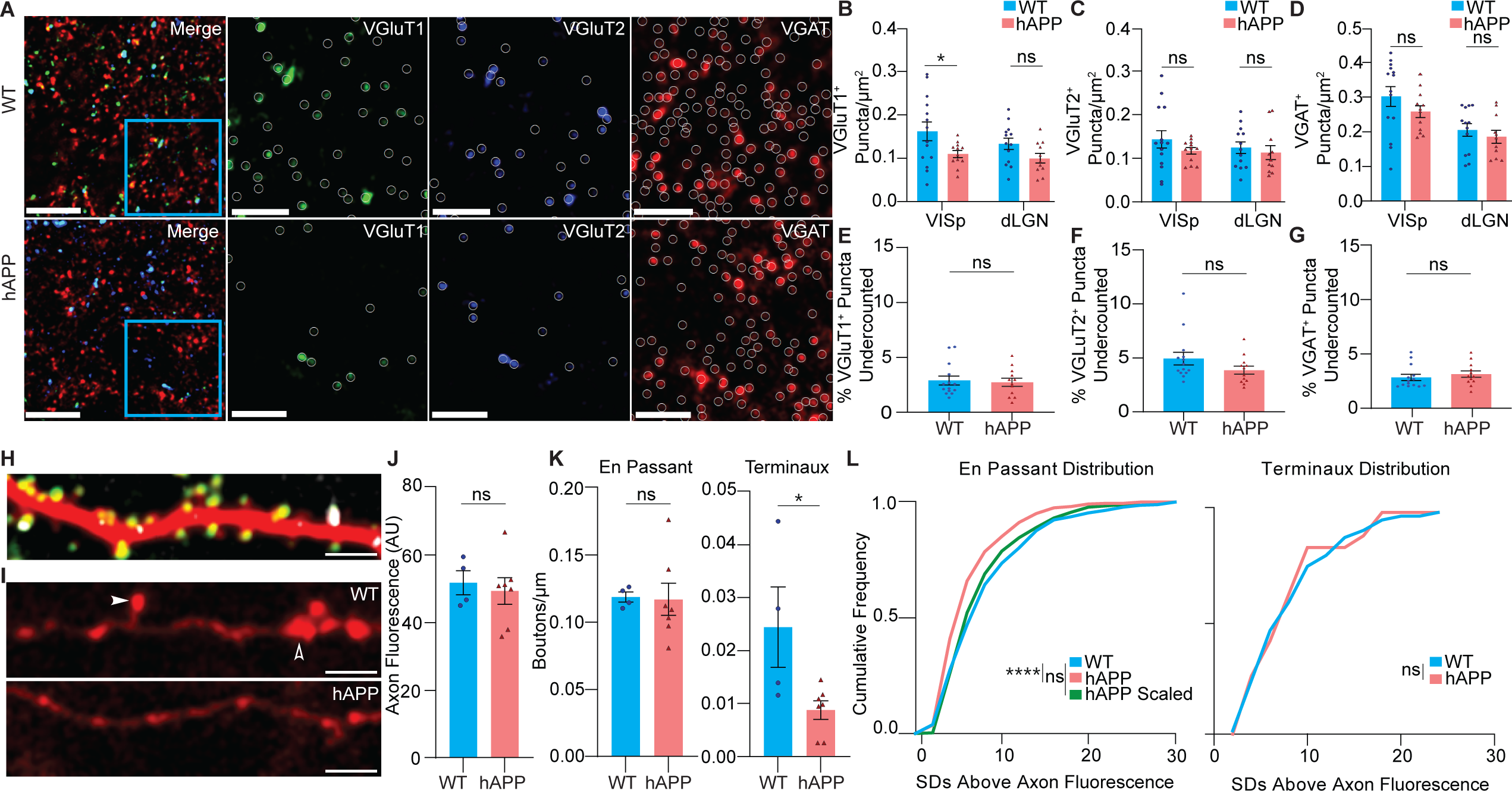
Synapse loss in the visual system is selective for cortical excitatory presynaptic terminals in hAPP mice. **A)** Representative images of synaptic immunohistochemistry labeling VGluT1^+^, VGluT2^+^, VGAT^+^, and merged puncta channels in layer 2/3 of the primary visual cortex in WT (upper) and hAPP (lower) mice housed under 12 h light/dark cycle. Circles indicate puncta identified by automated synapse detection. Some circles contain dim puncta that have sufficient fluorescence above the local background to meet the detection threshold. Scale bar: 10µm (merge), 5µm (individual channels). **B-D)** Density of VGluT1^+^ (B), VGluT2^+^ (C), and VGAT^+^ (D) puncta in the primary visual cortex (VISp) and dorsal part of the lateral geniculate nucleus (dLGN). ns, not significant (p > 0.05), *p < 0.05, unpaired student’s *t*-tests. n = 14 WT light, 12 hAPP light mice (VISp). n = 13 WT light, 11 hAPP light mice (dLGN). **E-G)** Estimated percentage of synaptic puncta undercounted in the visual cortex due to clustering in VGluT1 (E), VGluT2 (F), and VGAT channels (G). ns, not significant (p > 0.05), student’s *t*-test, n = 14 WT light, 12 hAPP light mice. **H)** Representative *in vivo* image of a pseudocolored dendrite (red) of an excitatory layer 2/3 primary visual cortical neuron with fluorescently labeled postsynaptic termini (PSD95, green, and gephyrin, white puncta). **I)** Representative WT axon (upper) and hAPP (lower) axons with example terminaux (filled arrowhead) and en passant (open arrowhead) boutons. Axons (I) in this dataset were differentiated from dendrites (H) by thin branch morphology and lack of fluorescently labeled postsynaptic structures. Scale bar for H-I: 5µm. **J)** Average fluorescence of axons on which boutons were analyzed. **K)** En passant (left) and terminaux (right) bouton density of excitatory layer 2/3 primary visual cortical neurons. Data are presented as mean ± SEM, and individual data points represent mouse averages. ns, not significant (p > 0.05), *p < 0.05, Mann-Whitney tests, n = 4 WT and 7 hAPP mice for J-K. **L)** Cumulative frequency distributions of imaging en passant (left) and terminaux (right) bouton brightness, represented as the number of standard deviations (SDs) above nearby axonal shaft fluorescence. WT and hAPP distributions are shown for both bouton types, and the hAPP distribution scaled by a factor of 1.22 is additionally shown for the en passant (left). ns, not significant (p > 0.05), ****p < 0.0001, Kolmogorov-Smirnov tests. n = 347 en passant and 62 terminaux boutons (WT), 287 en passant hAPP and 19 terminaux boutons (hAPP) analyzed from 4 WT and 7 hAPP mice.

### hAPP overexpression induces morphology-specific weakening and loss of excitatory presynaptic boutons *in vivo* but does not influence short-term dynamics

To further confirm that the reduction in VGluT1 in hAPP mice represents cortical excitatory synapse loss in the visual cortex, we analyzed the density and dynamics of presynaptic boutons *in vivo*. Datasets of excitatory layer 2/3 visual cortical neurons imaged from anesthetized head-fixed mice in two sessions separated by one week of standard 12 h light/dark housing using two-photon microscopy were used for these analyses (Niraula et al., 2023a; Niraula et al., 2023b). The neurons in this dataset were fluorescently labeled with TdTomato, a cell fill used to visualize dendritic and axonal morphology and excitatory (PSD-95-venus) and inhibitory (Teal-gephyrin) postsynaptic markers. Axons were identified by thinner shaft morphology and the absence of dendritic spines and labeled postsynaptic structures. We counted each en passant and terminaux presynaptic boutons (Figure 3I) whose brightness was greater than three standard deviations from the mean local axonal shaft fluorescence. We confirmed that the axonal shaft fluorescence was similar between WT-light and hAPP-light mice (Figure 3J). We found that en passant bouton density in hAPP mice was not different from that in WT mice (Figure 3K, left), but terminaux bouton density was significantly reduced (p<0.05, Mann-Whitney test, Figure 3K, right). However, the distribution of normalized fluorescence (standard deviations above the baseline) of en passant boutons was left-shifted in hAPP-light mice (p < 0.001, Kolmogorov-Smirnov test, Figure 3L, left), while terminaux bouton fluorescence did not differ (Figure 3L, right) compared to WT-light. We additionally found that the distribution of normalized en passant bouton fluorescence in WT-light mice is statistically indistinguishable from boutons from hAPP-light mice scaled by a factor of 1.22 (Figure 3K, left).

To test whether the structural dynamics of cortical excitatory presynaptic boutons are altered due to hAPP overexpression, we calculated the gain and loss of layer 2/3 visual cortical boutons in mice between two imaging sessions separated by one week of standard 12 h light/dark housing (Figure 4A). The percentage of stable, gained, and lost boutons were similar between the genotypes for both en passant and terminaux boutons (Figure 4B-D). Furthermore, en passant and terminaux boutons present in both imaging sessions showed no genotype-specific brightness changes between the sessions (Figure 4E).

**Figure 4:**
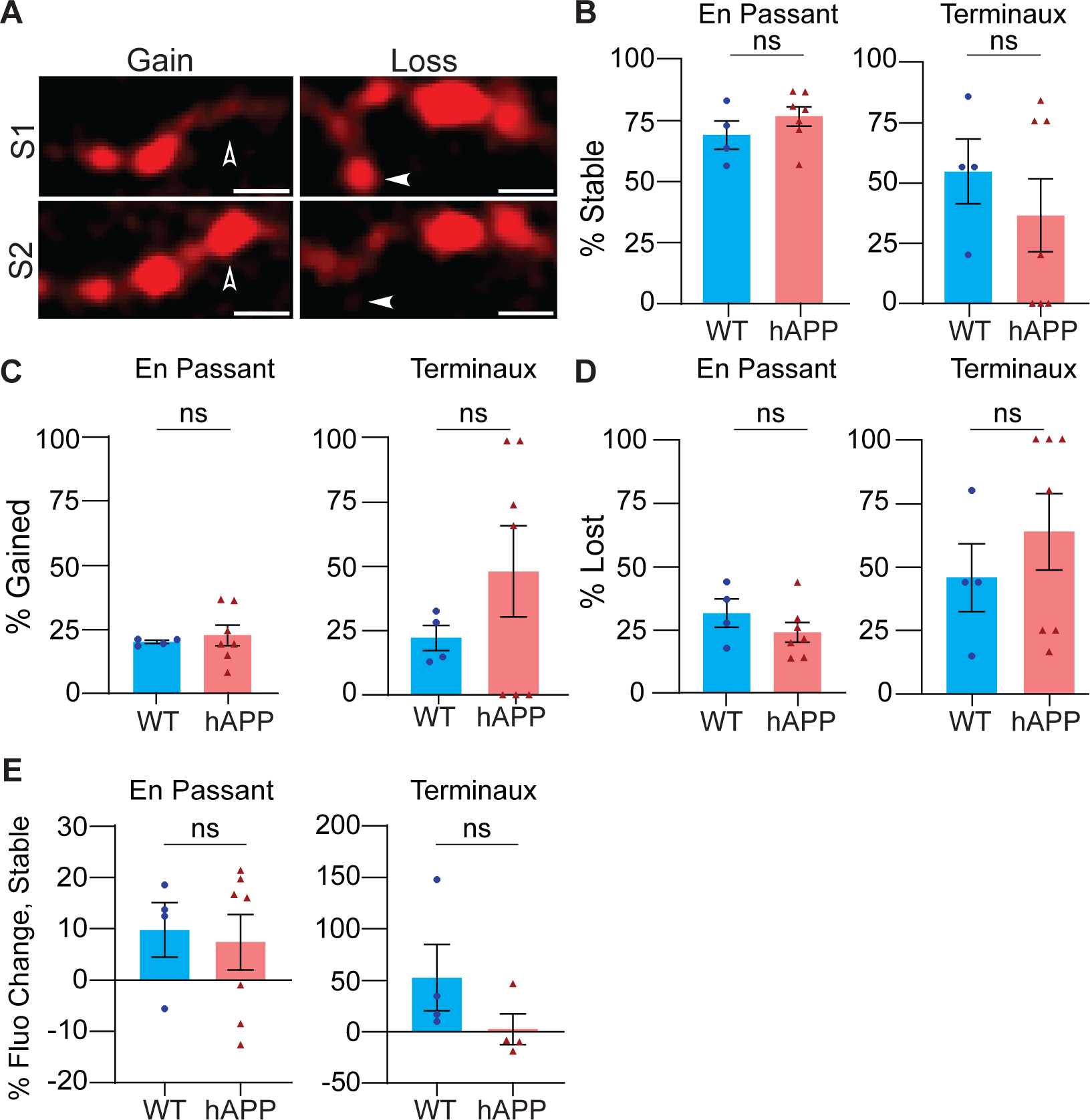
Excitatory layer 2/3 visual cortical bouton dynamics are unaltered across one week in hAPP mice *in vivo.* **A)** Representative images of boutons that are gained (left, open arrowhead) and lost (right, filled arrowhead) between imaging sessions. Scale bar: 2µm. **B-E)** En passant (left) and terminaux (right) bouton dynamics, including percent stable (B), gained (C), lost (D), and fluorescence change (E) of boutons in S2 relative to S1. Data are presented as mean ± SEM, and individual data points represent mouse averages. ns, not significant (p > 0.05), Mann-Whitney tests, n = 347 4 WT and 7 hAPP mice.

## Discussion

The primary goals of this study are to examine whether cellular and synaptic properties are differentially perturbed in cortical and subcortical regions and to identify whether visual network abnormalities are inherited from subcortical alterations or emerge locally in the cortex of hAPP mice. Functional connectivity in the brain networks, including the visual network, has been shown to contribute to the spread of amyloid and Tau in humans (Franzmeier et al., 2022; Franzmeier et al., 2020; Jagust and Mormino, 2011; Liu et al., 2023; Schoonhoven et al., 2023; Sintini et al., 2021). In this study, we show that functional connectivity significantly predicts aberrant expression of c-Fos, a critical gene for neural plasticity, in hAPP mice. Our results show that synaptic and cellular disruption are not inherited from subcortical regions but emerge in the visual cortex. However, within cortical areas, c-Fos abnormalities could emerge non-locally in brain regions due to their interconnectedness with affected areas. Functional connectivity only partly contributes to aberrant c-Fos expression; therefore, other factors not tested in the current study, such as the levels of oligomeric amyloid and local modulation of activity-dependent release of Aβ (Bero et al., 2011; Cirrito et al., 2005; Pignataro and Middei, 2017), may also contribute to differential c-Fos expression in brain regions.

The differences in cortical and subcortical regions of the visual network in hAPP mice are further underscored by differential but subtle alterations in presynaptic vesicular glutamate transporter VGluT1 and VGluT2 in these regions. The subtle presynaptic changes in the cortex resemble multiplicative synaptic scaling-like structural adaptations (Kim et al., 2012), as the normalized bouton fluorescence distribution in the visual cortex of this model mirrors a downscaled version of the WT distribution. The scaled distribution could reflect an adaptation to hAPP overexpression elicited aberrant activity, which could also contribute to elevated c-Fos expression in the cortex. Our observations indicate that cellular and synaptic abnormalities in the AD model represent a maladaptive transformation of the baseline physiological state seen in WT conditions rather than entirely sporadic and unlinked manifestations.

### Aberrant c-Fos expression and impaired experience-dependent modulation in the visual cortical regions of hAPP mice

c-Fos expression increases with a novel experience and is downregulated with habituation (Papa et al., 1993). Therefore, greater baseline c-Fos levels in hAPP mice in certain brain regions, including primary and higher-order visual cortical areas, could arise due to a defect in their ability to habituate to repeated everyday experiences. Consistently, AD patients and mouse models exhibit impaired suppression of neural activity elicited by repeated visual stimuli (Adams et al., 2021; Niraula et al., 2023a; Pihlajamaki et al., 2011). The inability to suppress irrelevant visual information could disrupt visual processing (Mielke et al., 1995) and lead to memory interference and cognitive deficits observed in AD (Drzezga et al., 2005; Niraula et al., 2023a; Pihlajamaki et al., 2011).

We expected that depriving light experience would reduce aberrant c-Fos expression since hAPP mice would receive no visual stimulation for their visual networks to interpret as novel. Interestingly, we found an increase in c-Fos expression after one week of light deprivation in WT mice, but no such increase was observed in the hAPP mice. The exception to this trend was a significant reduction of c-Fos expression in the superior colliculus, which receives direct retinal projections (Ito and Feldheim, 2018; Seabrook et al., 2017) of light deprived mice. In other regions, we speculate that prolonged darkness itself could represent novelty to the visual network, with non-visual or cross-modal inputs contributing more to its activity, leading to higher c-Fos expression in WT mice. Consistently, one week of light deprivation leads to the broadening of spontaneous but not visually evoked calcium transients in the visual cortex (Niraula et al., 2024). We speculate that the visual system of hAPP mice also senses the novelty of light deprivation and thus continues to express high c-Fos regardless of light experience. Though the effect of circadian rhythm shift in WT and hAPP mice (Kress et al., 2018) on c-Fos expression cannot be ruled out, the increase in c-Fos expression after one day of light deprivation and a lack of anxiety-like behavior after one week of light deprivation suggest that it may not play a major role. Future experiments with additional timepoint measurements of c-Fos expression are needed to confirm whether c-Fos levels decrease in WT mice as they habituate to extended light deprivation.

### Differential sensitivity of presynaptic termini subtypes in the visual network of hAPP mice

Similar to differences in aberrant c-Fos expression in cortical and subcortical regions, we also observed a subtle reduction in presynaptic puncta in the cortex but not the thalamus of the hAPP mice. Presynaptic termini are highly vulnerable at the initial stages of amyloid accumulation (Barthet and Mulle, 2020; de Wilde et al., 2016; Hark et al., 2021; He et al., 2019; Russell et al., 2012; Sze et al., 2000; Trujillo-Estrada et al., 2014). Reduced excitatory presynaptic density has repeatedly been shown to accompany amyloid accumulation (Bell et al., 2007; Canas et al., 2014; Kashani et al., 2008; Kirvell et al., 2006; Mitew et al., 2013; Mucke et al., 2000; Naito et al., 2017). Inhibitory synapses have also been shown to be vulnerable to amyloid toxicity, but the results are inconsistent (Agarwal et al., 2008; Canas et al., 2014; Hollnagel et al., 2019; Kurucu et al., 2022; Mitew et al., 2013; Naito et al., 2017; Niraula et al., 2023a; Palop et al., 2007; Ruiter et al., 2020; Verret et al., 2012).

We found VGluT1^+^ structures to be more vulnerable than VGluT2^+^ or VGAT structures in hAPP mice. Though we do not see amyloid plaque at this age, one possible cause could be higher Aβ levels in VGluT1^+^ boutons than in VGluT2^+^ boutons (Sokolow et al., 2012). An alternative possibility is that VGluT1^+^ puncta predominantly reflect presynaptic termini from cortical neurons, which display elevated c-Fos^+^ cell density, whereas VGluT2^+^ structures in the mammalian visual cortex represent projections of the dorsal lateral geniculate nucleus (Balaram et al., 2013; Balaram et al., 2011; Fremeau et al., 2001; Garcia-Marin et al., 2013; Nahmani and Erisir, 2005), which in contrast does not show significant changes to c-Fos^+^ cell density. VGluT1^+^ presynaptic loss may represent an adaptation to persistent novelty-like aberrant cortical activity in hAPP animals. However, further experiments are needed to test whether elevated c-Fos and presynaptic weakening are a consequence of aberrant activity and whether subcortical inputs expressing VGluT2 display less aberrant activity than VGluT1-expressing cortical inputs.

Interestingly, within excitatory synapses of cortical origin, we observed morphology-specific alterations in hAPP mice; en passant boutons are equally dense as WT mice, albeit with lower normalized fluorescence, whereas terminaux boutons are sparser in hAPP mice. Though functional differences between en passant and terminaux boutons are not well understood, morphology-based modeling suggests that boutons on extremely small terminal branches are especially sensitive to changes in membrane potential (Luscher and Shiner, 1990). Additionally, terminaux boutons are less likely than en passant boutons to harbor mitochondria, which are more commonly found in stable boutons than unstable boutons (Lees et al., 2019). These findings are also consistent with a recent study showing reduced terminaux bouton density in the barrel cortex of the same mouse model (Stephen et al., 2019). Overall, the presynaptic phenotypes are subtle at this age in the visual cortex of hAPP mice, potentially due to an intrinsic property of the visual cortex, lack of amyloid accumulation, or lack of Tau pathology in our model. Additionally, the limited number of mice per sex per group precluded discerning any sex effects. The subtle synaptic changes compared to changes in c-Fos expression and functional connectivity demonstrates that network changes may precede large-scale synaptic changes in the visual system. Therefore, the absence of overt neuropathological changes in the visual system of early-stage AD patient brains need not imply a lack of functional plasticity disruption.

## Materials and Methods

### Mice

All animal procedures are approved by the University of Kansas Institute of Animal Use and Care Committee and meet the NIH guidelines for the use and care of vertebrate animals. J20 mice (B6.Cg-Tg(APPSwFlLon, PSEN1*M146L*L286V)6799Vas/Mmjax, RRID: MMRRC_034848-JAX) was originally obtained from the Mutant Mouse Resource and Research Center (MMRRC) at The Jackson Laboratory, an NIH-funded strain repository, and was donated to the MMRRC by Robert Vassar, Ph.D., Northwestern University. They were maintained as heterozygotes for the hAPP transgene by breeding heterozygous J20-hAPP male mice with 1-2 C57BL/6J female mice. Litters from these matings consisting of both male and female heterozygous J20-hAPP mice and their non-transgenic C57BL/6J background (WT) littermates were used for experiments. A maximum of five mice were housed in a standard cage but individually housed after the cranial window surgery. Mice were housed on a 12 h light/12 h dark cycle except for the group that went through a period of light deprivation (24 h dark). For light deprivation, mice were placed in a ventilated, lightproof cabinet inside of a dark room for seven days. Infrared goggles with a low intensity 850 nm wavelength emission source were used at all times while maintaining mice in dark housing to prevent light exposure. All mice used for c-Fos immunohistochemistry were housed with 2-3 littermates. For c-Fos immunohistochemistry, functional connectivity, and plasticity deficit index analyses, n = 8 WT-light (5 males, 3 females), 9 WT-dark (4 males, 5 females), 10 hAPP-light (4 males, 6 females), and 8 hAPP-dark mice (3 males, 5 females) aged 4-6 months were used. For c-Fos/NeuN colocalization analysis, n = 4 WT-light, 5 hAPP-light male mice aged 5 months housed in a standard 12 h light/dark cycle were used. For circadian behavior analysis, n = 12 WT mice (6 males, 6 females) aged 4-6 months were used. Half these mice (n = 6 WT mice [3 males, 3 females]) underwent modified open field testing after one week of 12 h light/dark cycle, and the other half underwent modified open field testing after one week of 24 h darkness. For widefield presynaptic immunohistochemistry, n = 14 WT-light (9 males, 5 females) and 12 hAPP-light mice (6 males, 6 females) aged 4-6 months were used. For high-resolution confocal presynaptic imaging and analysis, n = 12 WT-light (7 males, 5 females) and 11 hAPP-light mice (5 males, 6 females) randomly selected from the widefield presynaptic immunohistochemistry group were reimaged.

### Circadian and Anxiety-Related Behavior

To analyze shifts in circadian- and anxiety-related behavior following one week of visual deprivation, 4–6-month-old WT mice sorted into four cages of three mice each were used for both circadian behavior and modified open field experiments. The mice were housed under a 12-hour light/dark cycle for one week, followed by 24-hour visual deprivation for one week. To assess circadian shifts, cages were monitored throughout the duration of the paradigm using an infrared surveillance system. Mice were individually tracked in each cage based on the placement of ear tags. The videos were binned into 30-minute segments, and each individual mouse was recorded as “active” for each bin if it moved entirely out of the nest at any point. Mouse activity was analyzed for the first and last days of the standard 12 h light/dark and 24 h dark cycle weeks.

Half of the mice underwent an open field test day after one week of a 12 h light/dark cycle, and the remaining half underwent an open field test day after one week of 24 h light deprivation. The open field consisted of placing each mouse in a 40cm × 40cm test chamber equipped with a force plate actometer beneath the chamber’s floor to track the mouse’s position. The first five minutes of the test were conducted in total darkness (Dark OF) and the last five minutes were conducted with the experimental room light on (Lit OF). The amount of time that each mouse spent exploring the center of the chamber was calculated by dividing the total time each mouse spent moving at over 5 cm/s in the center 80% of the chamber, by the total time each mouse spent moving at over 5 cm/s. The dark phase of the test was conducted before the light phase to prevent the mice that had undergone one week of dark housing from potentially displaying sudden light exposure-related anxious behavior rather than anxious behavior resulting from dark housing. Open field tests were all conducted during the light portion of the standard 12 h light/dark cycle for both groups.

### Tissue Preparation and Immunohistochemistry

Mouse cages were brought to the surgical suite and remained undisturbed for at least 5 h before brain extractions to avoid capturing c-Fos expression elicited by cage movement or contextual novelty. Additionally, all transcardial perfusions were performed near the center of the light phase of the standard 12 h light/dark cycle to avoid circadian transition-related c-Fos expression. Dark-housed mice were brought to the surgical suite in a light-blocking container and anesthetized in a dark room using infrared goggles to prevent light-evoked c-Fos expression. 4-6-month-old (presynaptic terminal immunohistochemistry) or 5-6-month-old (c-Fos immunohistochemistry) J20-hAPP and WT littermate mice were deeply anesthetized by intraperitoneal injection of 2% avertin in phosphate-buffered saline (PBS), pH 7.4 and transcardially perfused with cold PBS followed by 4% paraformaldehyde. The brains were extracted and post-fixed in 4% PFA overnight at 4°C, followed by storage in PBS. For 40 μm vibratome sectioning, the brains were embedded in 4% oxidized agarose as previously described (Ragan et al., 2012) to limit artifacts during sectioning on a vibratome (Leica VT1000 S). For 20 μm cryotome sectioning, brains were cryoprotected overnight at 4°C in 15% (w/v) and then in 30% (w/v) sucrose in phosphate buffer (PB). The brains were sectioned coronally on a microtome (Leica SM 2010R) and collected in PBS with sodium azide (0.02%).

24 evenly spaced 40 μm sections from each brain spanning the posterior midbrain to the anterior olfactory bulb were fluorescently immunolabeled for c-Fos, and 3 evenly spaced 20-40 μm sections spanning the visual cortex and dorsal part of the lateral geniculate nucleus (1-2 sections for each region per mouse) were fluorescently immunolabeled for VGAT^+^, VGluT1^+^, and VGluT2^+^ puncta. Sections were permeabilized for 2 h at room temperature in a 1% TritonX-100 and 10% normal goat serum (NGS) solution in PBS followed by incubation with mouse anti-NeuN (1:250, Sigma cat. #MAB377) and/or rabbit anti-c-Fos (1:1000, CST cat. #2250S) or rabbit anti-VGAT (1:1000, Synaptic Systems cat. #131002), mouse anti-VGluT1 (1:2000, Sigma cat. #MAB5502), and guinea pig anti-VGluT2 (1:1000, Sigma cat. #AB2251-I) in a PBS solution containing 0.1% TritonX-100 and 5% NGS overnight at 4°C. Sections were then washed 3× with PBS and incubated with Alexa 647-conjugated goat anti-mouse antibody (1:2000, Fisher cat. #A21236) for NeuN immunohistochemistry and/or Alexa 555-conjugated goat anti-rabbit antibody (1:2000; Fisher cat. #PIA32732) for c-Fos immunohistochemistry or Alexa 488 conjugated goat anti-rabbit antibody (1:2000, Fisher cat. #A11008), Alexa 555-conjugated goat anti-mouse antibody (1:2000, Fisher cat. #A32727), and Alexa 647-conjugated goat anti-guinea pig antibody (1:2000, Fisher cat. #A21450) for presynaptic terminal immunohistochemistry for 2 h in a PBS solution containing 0.1% TritonX-100 and 5% NGS at room temperature, followed by three washes with PBS before mounting on glass slides.

### *In Vivo* Presynaptic Bouton Structural Labeling

Images used for in vivo presynaptic structural analysis were previously acquired as described (Niraula et al., 2023a; Niraula et al., 2024), whereas all other experiments presented were generated specifically for this study. Previously published analyses of the same in vivo experimental images were focused on analyzing fluorescently labeled postsynaptic PSD-95 and gephyrin puncta on dendritic spines and the shaft. In this study, only presynaptic boutons on axons labeled with TdTomato cell fill were independently analyzed (Figures 3H-L and 4). Briefly, plasmids expressing TdTomato cell fill (to label the cell and visualize morphology, including presynaptic structures), Teal-gephyrin (to visualize inhibitory postsynaptic termini), and PSD95-venus (to label excitatory postsynaptic termini) in a Cre recombinase (Cre) dependent manner along with a plasmid expressing Cre were injected into the right lateral ventricle of embryonic days 15.5-16.5 J20 heterozygote and WT littermate embryos in utero to fluorescently label excitatory L2/3 neurons as previously described (Subramanian et al., 2019; Tabata and Nakajima, 2001; Villa et al., 2016). To achieve sparse labeling of neurons with all fluorophores for uncluttered visualization, the Cre-dependent plasmids expressing the fluorescent markers were injected at high concentrations to increase the likelihood of all three plasmids being taken up by the same neuron, while the plasmid expressing the Cre recombinase was injected at a low concentration so that a sparse set of cells would express the fluorophores. A pair of platinum electrodes (Protech International) and a square wave electroporator (ECM830, Harvard Apparatus) were used to target these plasmids toward the visual cortex. Pups born ∼3-4 days after electroporation were implanted with a cranial window after reaching 4-6-months of age. Briefly, a 5 mm circle of the skull was removed and replaced with a glass coverslip over the visual cortex of the right hemisphere, and a titanium headpost for head-fixed imaging was adhered to the skull using dental cement. Optical intrinsic signal imaging was performed 14 days later to map the location of the visual cortex (Kalatsky and Stryker, 2003; Niraula et al., 2023a). L2/3 excitatory visual cortical neurons were imaged in anesthetized head-fixed mice in two sessions separated by one week of normal 12 h light/dark cycle.

### Image Acquisition

For widefield imaging, brain sections immunostained for c-Fos or presynaptic markers were imaged using an ImageXpress Pico automated imaging system (Molecular Devices, San Jose, CA) with a 10× objective (c-Fos; Leica HC PL FLUOTAR 10×/0.32) or a 63× objective (presynaptic markers; Leica HC PL FLUOTAR 63×/0.70). High-resolution images of presynaptic puncta were acquired using a Nikon AX confocal microscope equipped with GaAsP detectors and a Nikon Plan APO Lambda D 60×/1.42 oil-immersion objective and 4× zoom for a total magnification of 240×. Images used for *in vivo* presynaptic structural analysis were acquired using a two-photon microscope (Sutter Instruments) equipped with Nikon 16x/0.8 NA objective and GaAsP detectors (Niraula et al., 2023a; Niraula et al., 2024).

### Data and Statistical Analysis

Brain section mapping into a common mouse brain atlas coordinate framework, cell/synapse detection, puncta fluorescence collection, and brain region area measurements were performed using NeuroInfo software (MBF Bioscience, Williston, VT). 12-bit images of brain sections were first mapped in 3D via both linear and nonlinear splining to the Allen CCFv3 atlas space to allow for automated cell/synapse detection and area measurement by region. A total of 31 regions covering the entire cerebral cortex, cerebral nuclei, interbrain, and midbrain, as well as 17 subregions of the visual network were mapped for analysis. Bright circular objects against a darker background were automatically detected using a scale-space analysis of the response to Laplacian of Gaussian (LoG) within the expected diameter range (8-18 µm) of labeled cell body or presynaptic puncta diameters as described (Kong et al., 2013). Briefly, pixel fluorescence values were filtered with an omnidirectional LoG transformation to identify circular objects with fluorescence above the local background. Next, above-threshold groups of pixels within an expected range of cell body (8 - 18 µm) or presynaptic terminal diameters (0.32-1.97 µm) were identified as potential objects. A user-defined LoG threshold was then applied to discriminate cells/synapses from false positives identified due to background fluorescence fluctuations. Only objects above the respective LoG strength threshold were included in the analysis, and identified objects were manually proofread to eliminate remaining false positives. The c-Fos cell detection threshold was set such that a large number of dimly labeled cells that represent background or near-background c-Fos expression or false positives from background fluorescence fluctuation (∼70% of detected “objects” with expected nucleus size criteria) were not counted as “c-Fos^+^.” A histogram showing the number of detected objects at each LoG threshold value was used to assist with determining the LoG values below which false positives would increasingly be identified. The synaptic punctum detection threshold was set in the same manner: the lowest possible LoG threshold showing minimal false positives was identified for each channel. Cell/synaptic punctum detection thresholds were set based on these criteria for WT mice, then applied to all images of the same channel and imaging conditions uniformly for both genotypes. Based on these criteria, the LoG threshold value was set at LoG threshold = 55 for cellular nuclei and 101 for all presynaptic puncta captured through lower-resolution light microscopy and 2D puncta detection (range 0-255). For identifying puncta from 3D reconstruction z-stacks captured via high-resolution confocal imaging, a volumetric LoG filter was applied in 3D through the z-stack with thresholds of 101 and 56, respectively, for VGAT and VGluT2 puncta. Small VGluT1 puncta rapidly photobleached during high-resolution confocal microscopy and, therefore, were not analyzed. All detected synaptic and cellular puncta were manually proofread for accuracy following automated detection.

For c-Fos/NeuN colocalization analysis, the primary visual cortex from three single-hemisphere sections spanning the posterior to the anterior primary visual cortex was analyzed per mouse. NeuN^+^ cells were detected using a LoG = 43 filter and were automatically notated as colocalized with c-Fos^+^ cells (LoG = 55 filter) if the markers identifying the center of each respective object were within half of an expected nucleus diameter of each other (within 6.5 µm). All identified c-Fos^+^ cells were then manually proofread for accuracy for colocalization or lack of colocalization with NeuN signal from a cell of the same shape and position. The fraction of c-Fos^+^ cells colocalized with NeuN^+^ cells was calculated by dividing the number of c-Fos^+^+NeuN^+^ cells by the number of c-Fos^+^ cells for each mouse and compared using a Mann-Whitney test.

Cell/synaptic puncta density for each mouse was calculated by dividing the total number of cells or synapses per region by the area (or analyzed volume for z-stacks) per region across all sections for each brain (24 sections for c-Fos, 2-3 sections for synapses). All regions in both hemispheres of the 24 brain-wide sections were analyzed for c-Fos^+^ cell detection. A 3-way mixed-effects restricted maximum likelihood ANOVA was performed for each network to determine the effects of genotype (between-subject), light experience (between-subject), brain region (within-subject), and their interactions on c-Fos^+^ cell density. Since both networks showed significant interactions between all combinations of factors, 2-way ANOVAs (genotype [between-subject], light experience [between-subject], and their interaction) were repeated for each brain region to determine which specific brain regions show significant interaction following Benjamini-Hochberg false discovery rate correction. Significant interactions were then used for post-hoc Šídák’s multiple comparisons test (1 family with 4 comparisons: WT-light vs hAPP-light, WT-dark vs. hAPP-dark, WT-light vs. WT-dark, hAPP-light vs. hAPP-dark).

For open field analysis, a 2-way mixed effects restricted maximum likelihood ANOVA (light experience [between-subject], open field light/dark [within-subject] interaction) was performed when the mouse moves at a speed >5 cm/s in the center 80% of the chamber. For functional connectivity analysis, Pearson’s *r* was calculated between each pairwise combination of brain regions either within the whole brain or visual network for light and dark conditions, and average z values were calculated following Fisher’s r-to-z transformation for each region within its respective network. For each region, the effect size of hAPP on c-Fos^+^ cell density was calculated as:

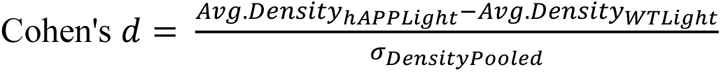

For each region, the c-Fos modulation deficiency index *i* was calculated as the absolute value of the effect size of hAPPDark/hAPPLight – WTDark/WTLight c-Fos^+^ cell density values:

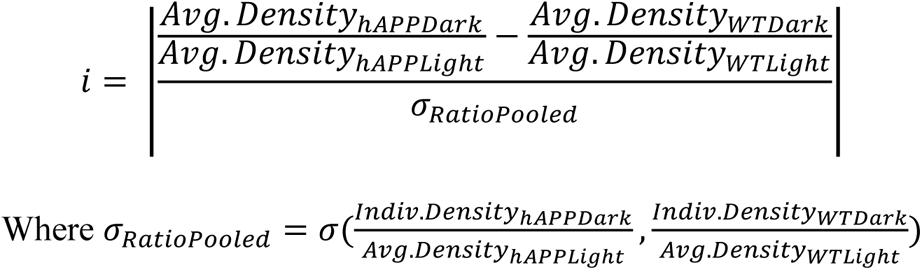

or the pooled standard deviation of each individual hAPP mouse’s hAPP-dark density divided by the average hAPP-light and each individual WT mouse’s WT-dark density divided by the average WT-light density and density combined.

For widefield and high-resolution presynaptic immunohistochemistry density comparisons and for synaptic puncta clustering comparisons, unpaired *t-*tests were run on each synapse subtype individually. For *in vivo* presynaptic bouton density, axon fluorescence, and bouton dynamic data, Mann-Whitney tests were performed separately for each comparison. For *in vivo* bouton fluorescence cumulative distribution comparisons, Kolmogorov-Smirnov tests were performed.

For presynaptic terminal quantification from widefield 2D images, 1-2 ∼230 × 900 μm columns per mouse spanning all layers of one hemisphere of the visual cortex was analyzed, and 1-2 ∼230 × 375 μm columns per mouse vertically spanning one hemisphere of the dorsal lateral geniculate nucleus were analyzed. Localized areas containing artifacts were excluded from the analysis. Synaptic puncta fluorescence analysis in the visual cortex included a total of 759,230 WT-light and 550,683 hAPP-light VGAT^+^ puncta, 386,322 WT-light, and 223,032 hAPP-light VGluT1^+^ puncta, and 330,103 WT-light and 239,168 hAPP-light VGluT2^+^ puncta. For presynaptic terminal clustering percentage evaluation, a 10µm × 900µm column for each mouse spanning all layers of one hemisphere of the visual cortex was analyzed. All detected puncta in the columns were evaluated for incorrect clustering, and initially, uncounted puncta were added. The fraction of undercounted puncta was calculated by dividing the number of uncounted puncta by the number of automatically detected puncta.

For visual cortex presynaptic terminal quantification from confocal high-resolution images, three z-stacks of 73.5µm × 73.5µm × 0.18µm planes (∼10 z-planes per stack, average volume of 9,347.65µm^3^ per stack) were taken, one stack each from upper, middle, and lower layers of a single hemisphere of the primary visual cortex. Six z-stacks from two sections were quantified per mouse. High-resolution puncta density analysis included a total of 308,544 WT-light and 276,364 hAPP-light VGAT^+^ puncta and 375,508 WT-light and 368,876 hAPP-light VGluT2^+^ puncta.

For analysis of *in-vivo* presynaptic boutons, TdTomato-filled axons were first manually identified based on thinner branch morphology and absence of postsynaptic markers (PSD-95 and Teal-gephyrin), then en passant (centered on axonal shafts) and terminaux boutons (protrusions from axonal shafts) were manually labeled based on morphology using a modified version of the ObjectJ plugin (Villa et al., 2016) for FIJI (Schindelin et al., 2012). Puncta on or protruding from axonal shafts were scored as boutons only if they were present in two consecutive z-frames. After another investigator proofread the bouton identification and classification in both sessions, the marked boutons were subjected to an unbiased intensity-based threshold relative to the nearby axonal shaft. 5-pixel × 5-pixel measurement ROIs were placed over the identified boutons and along their respective axonal shafts, and the fluorescent intensity of each bouton and its nearest 10 neighboring axonal shaft ROIs were collected. Using the modified ObjectJ plugin, bouton IDs and locations were transferred from S1 to S2 image stacks, and newly formed and lost boutons were manually identified. Identified objects were excluded if their fluorescence did not exceed three times the standard deviation of fluorescence of the nearest neighboring axonal shaft ROIs. Bouton gain and loss were calculated as the number of lost or gained boutons between imaging sessions 1 (S1) and 2 (S2) divided by the number of boutons present in S1(for lost) or S2 (for gained). The percentage of stable boutons was calculated as the number of boutons that were present in both S1 and S2 divided by the number of boutons in S1. The percentage of fluorescence change in stable boutons between S1 and S2 was calculated as S2 fluorescence divided by S1 fluorescence for each stable bouton where a bouton’s fluorescence is represented by the number of standard deviations above the average fluorescence of the nearest 10 neighboring shaft ROIs in its respective session. Calculations for *in vivo* bouton data were performed using custom MATLAB scripts (https://github.com/JaiSubKU/bouton_analysis). For S1 bouton fluorescence distribution, a total of 347 en passant and 62 terminaux boutons were identified from WT-light, and 287 en passant and 19 terminaux boutons were identified from hAPP-light mice. Data normality for statistical analyses was assessed by Shapiro-Wilk tests. All statistical tests were performed in GraphPad Prism (GraphPad Software, San Diego, CA).

## Author contributions

OJL, ASS, and JS designed the experiments and analysis. OJL, JM, SN, and GD performed the experiments and analyzed the data. OJL and JS wrote the manuscript. AAS and SSY performed high-resolution confocal images, JS supervised the research.

## Acknowledgments

We thank the Subramanian lab members for their help with revising this manuscript, along with Connor Maher for his help with circadian-related behavior scoring. This work was supported by a grant from the National Institutes of Health (Grant No. RO1AG064067) to JS. JM was funded by a grant from the National Institute of General Medical Sciences of the National Institutes of Health: T34GM136453.

---

**Supplementary Figure 1:**
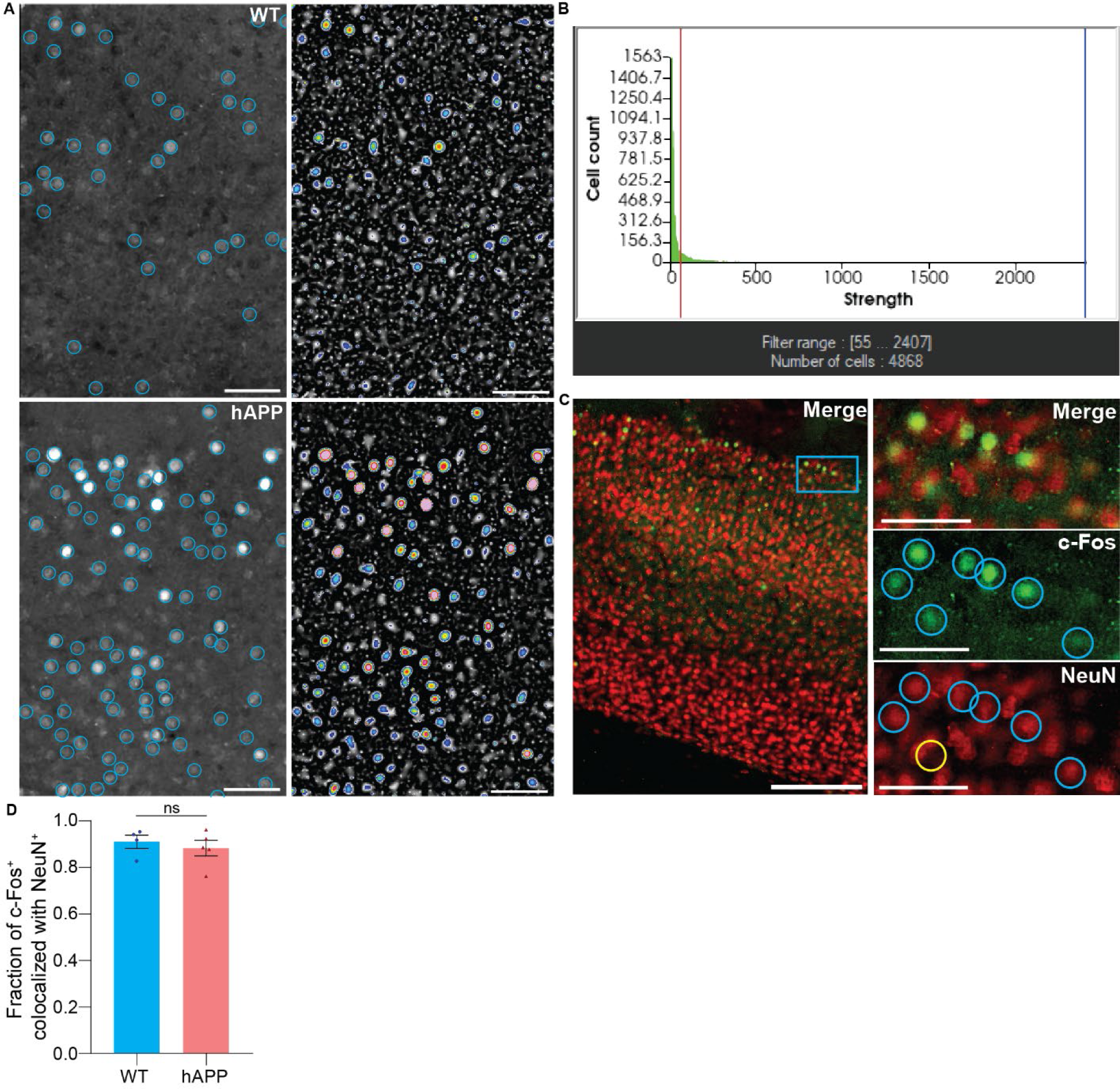
Visualization of c-Fos^+^ cell detection and colocalization with neuronal marker NeuN in the primary visual cortex. **A)** Representative images showing cell detection output (left panel) and the Laplacian of Gaussian (LoG) filtered images of the same field of view (right panel) in the WT (upper) and hAPP (lower) primary visual cortex. In each LoG image, pixels under the detection threshold (LoG = 55) are shown in greyscale, and pixels above the detection threshold are shown in a 14-color heatmap with warmer colors representing higher LoG values. Scale bar = 100µm. **B)** Histogram showing number of “objects” meeting the c-Fos^+^ nucleus size criteria (8-18µm) detected at each LoG (Strength) threshold in a full slice of the WT mouse shown in A). The LoG threshold for c-Fos detection was set at the minimum LoG threshold in WT animals, below which cells with near-background c-Fos expression or false positives from background fluctuation (∼70% of “objects”) are detected, and applied uniformly to all slices of both genotypes. Only “objects” above LoG = 55 were considered as “c-Fos^+^” cells. The red line indicates LoG = 55. The x-axis spans the full range of the data, though some data points with high LoG values are not visible due to the large number of detected “objects” at the lower end. **C)** Representative images showing merged (left) and individual c-Fos and NeuN channels in the WT primary visual cortex (right). The blue box is shown in higher magnification in panels to the right. The location c-Fos^+^ cells identified by the LoG = 55 threshold are shown in blue circles on the c-Fos channel and overlaid onto the NeuN channel. The yellow circle indicates a detected c-Fos^+^ cell without NeuN colocalization. Scale bar = 200µm for left merged, scale bar = 50µm for all others. Note that some cells show brighter c-Fos signal than NeuN signal or vice versa**. D)** The fraction of identified c-Fos^+^ cells colocalizing with NeuN in the primary visual cortex for WT and hAPP mice under a 12h light/dark cycle. Perfusions were performed close to the center of the light phase in the standard 12h light/dark cycle to avoid circadian transition-related effects. Data are presented as mean ± SEM with each data point representing an individual mouse. ns, not significant (p > 0.05), Mann-Whitney test, n = 4 WT and 5 hAPP mice, 3 slices spanning the primary visual cortex for each.

**Supplementary Figure 2:**
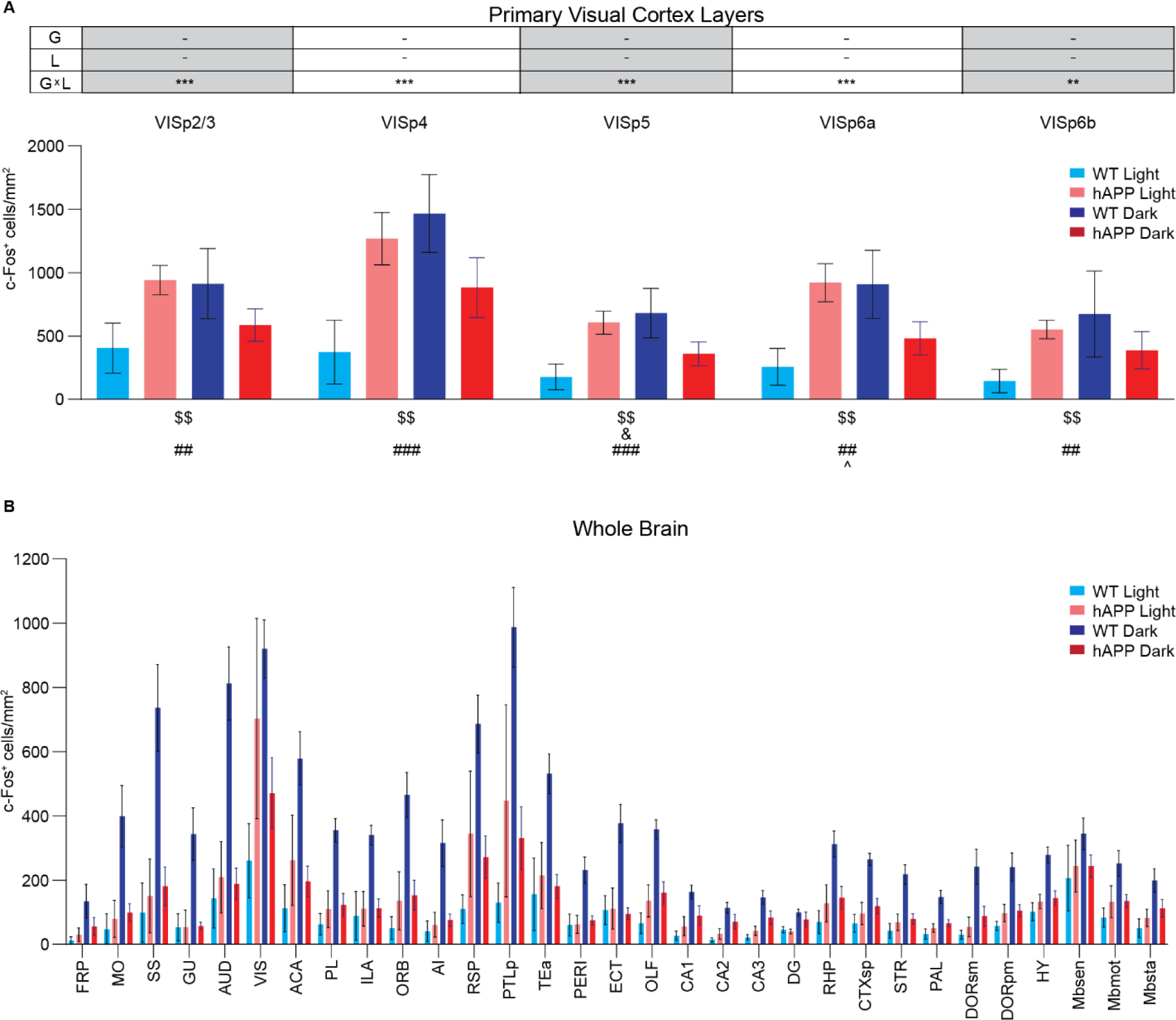
Extended data: hAPP overexpression or light deprivation increases c-Fos^+^ cell density. **A)** 2-way ANOVAs (WT/hAPP [G], Light/Dark [L], interaction [G × L]), with Benjamini-Hochberg’s false discovery rate correction for brain regions shown above. Post-hoc comparisons (Šídák’s multiple comparisons test, 1 family with 4 comparisons) are shown for regions with a significant interaction. F(1, 31) for all factors in all ANOVAs. Benjamini-Hochberg corrected p-values: -p > 0.05, **p < 0.01, ***p < 0.001. Post-hoc comparisons are shown below. WT Light vs. hAPP Light: ^$$^p < 0.01, WT Dark vs. hAPP Dark: ^&^p < 0.05, WT Light vs. WT Dark: ^##^p < 0.01, ^###^p < 0.001, hAPP Light vs. hAPP Dark: ^p < 0.05. **B)** c-Fos^+^ cell density for regions across the brain. See Table 1 for results of 2-way ANOVAs with post-hoc comparisons for regions with significant interaction. Data are presented as mean ± SEM. Perfusions for both Light and Dark groups were performed close to the center of the light phase in the standard 12h light/dark cycle to avoid circadian transition-related effects. n = 8 WT light, 9 WT dark, 10 hAPP light, and 8 hAPP dark housed mice.

**Supplementary Figure 3:**
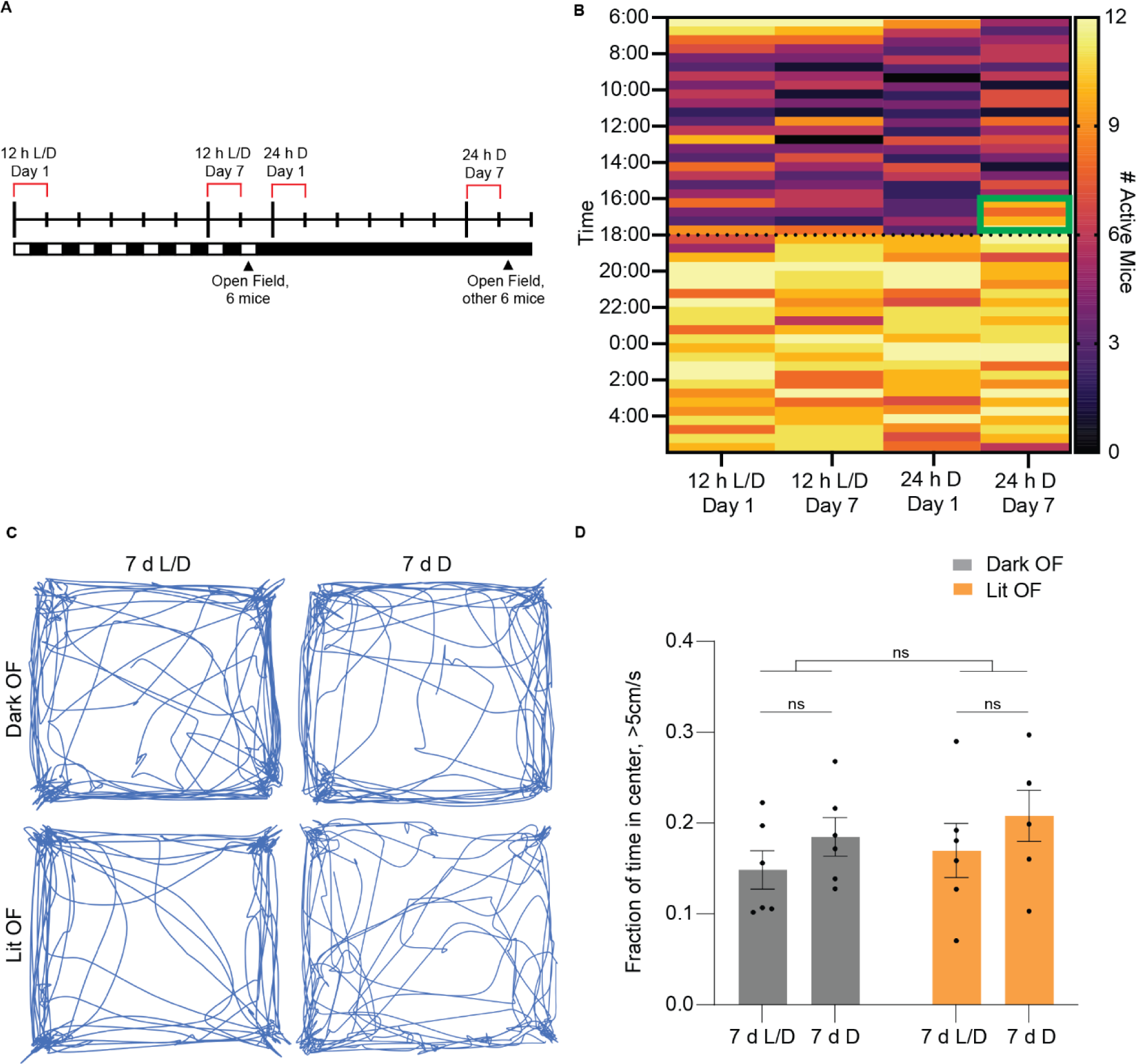
One week of light deprivation induces mild circadian drift but does not increase behavioral anxiety. **A)** Timeline for capturing circadian and anxiety-related behavior under a 12 h light/dark cycle (left half) and under 24 h darkness (right half). 4 cages of 3 mice each were monitored with an infrared surveillance system for 7 d of a 12 h light/dark cycle followed by 7 d of 24 h light deprivation. Each individual mouse was manually scored as “active” or “inactive” in 30-minute bins spanning 12 h L/D day 1, 12 h L/D day 7, 24 h D day 1, and 24 h D day 7 (red brackets). Half of the mice underwent an open field test after seven days of 12 h L/D, and the other half underwent the open field test after seven days of light deprivation (arrows). **B)** Heatmap of the number of mice active during each 30m interval on the 1^st^ day of 12 h L/D, the 7^th^ day of 12 h L/D, the 1^st^ day of 24 h D, and the 7^th^ day of 24 h D. The green box indicates increased mouse activity ∼2h before the typical light to dark phase transition time (18:00) in 24 h D Day 7 mice in contrast to other groups. **C)** Representative traces of mouse position in the dark (upper) or lit (lower) half of the open field test following one week of 12 h L/D (left) or 24 h D (right). **D)** Fraction of time spent moving over 5cm/s in the center 80% of the open field chamber in the first 5 m (Dark OF) and last 5 m (Lit OF) of the modified open field test following either 7 d of 12 h L/D or 7 d of 24 h D. ns, not significant (p > 0.05) for all comparisons, 2-way mixed effects restricted maximum likelihood ANOVA with repeated measures (Dark/Lit OF, Dark/Light housing, interaction, mouse matching, F[1,10] for between factors), n = 6 WT mice tested after 7 d L/D, separate 6 WT mice tested after 7 d D.

**Supplementary Figure 4:**
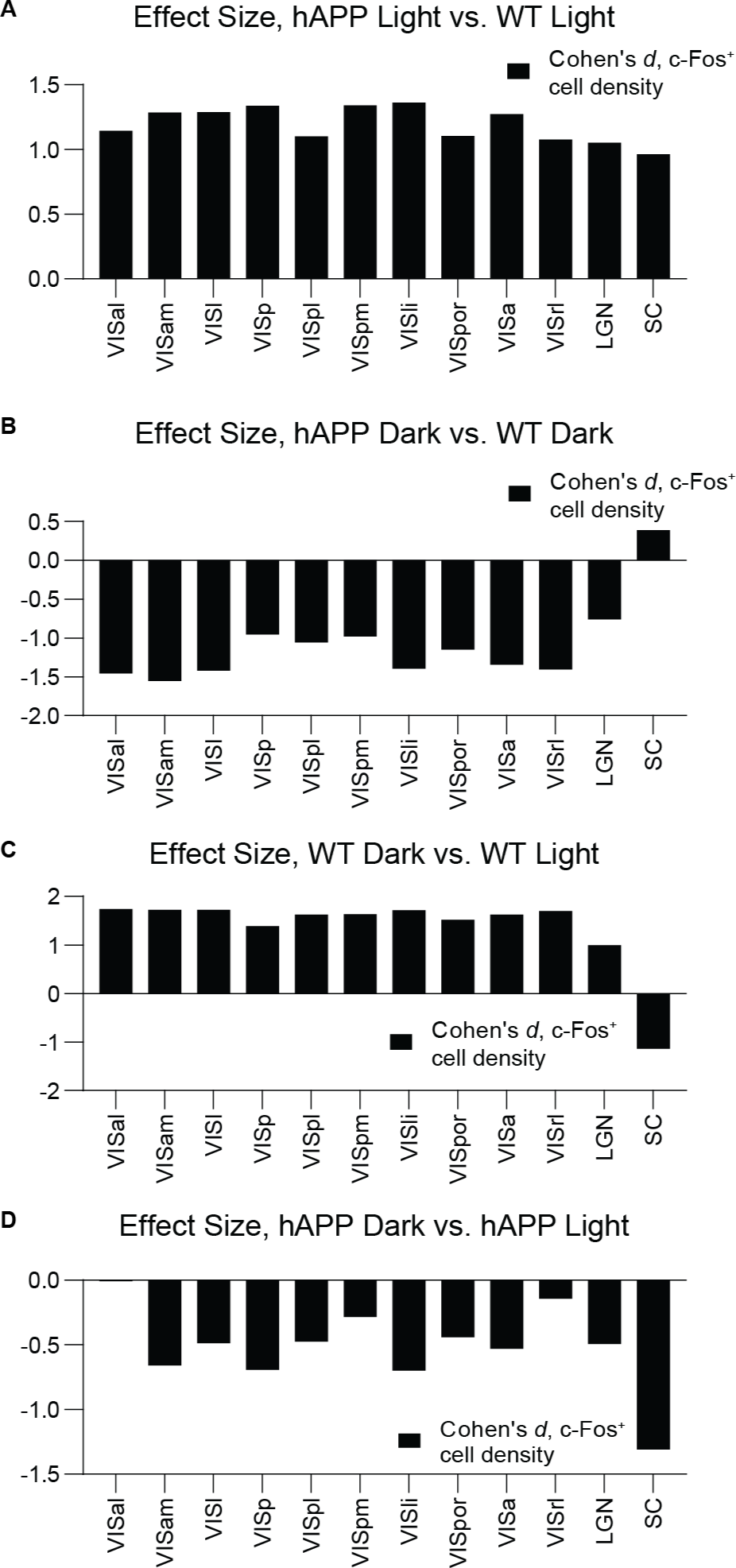
Effect size of hAPP overexpression and light deprivation on c-Fos^+^ cell density in the visual network. **A-B)** Effect size (Cohen’s *d*) of hAPP on c-Fos^+^ cell density for mice housed for one week under a normal 12h light/dark cycle (A) or 24h dark (B). **C-D)** Effect size of one week of 24h dark housing on WT (C) and hAPP mice (D). n = 8 WT light, 9 WT dark, 10 hAPP light, and 8 hAPP dark housed mice.

**Supplementary Figure 5:**
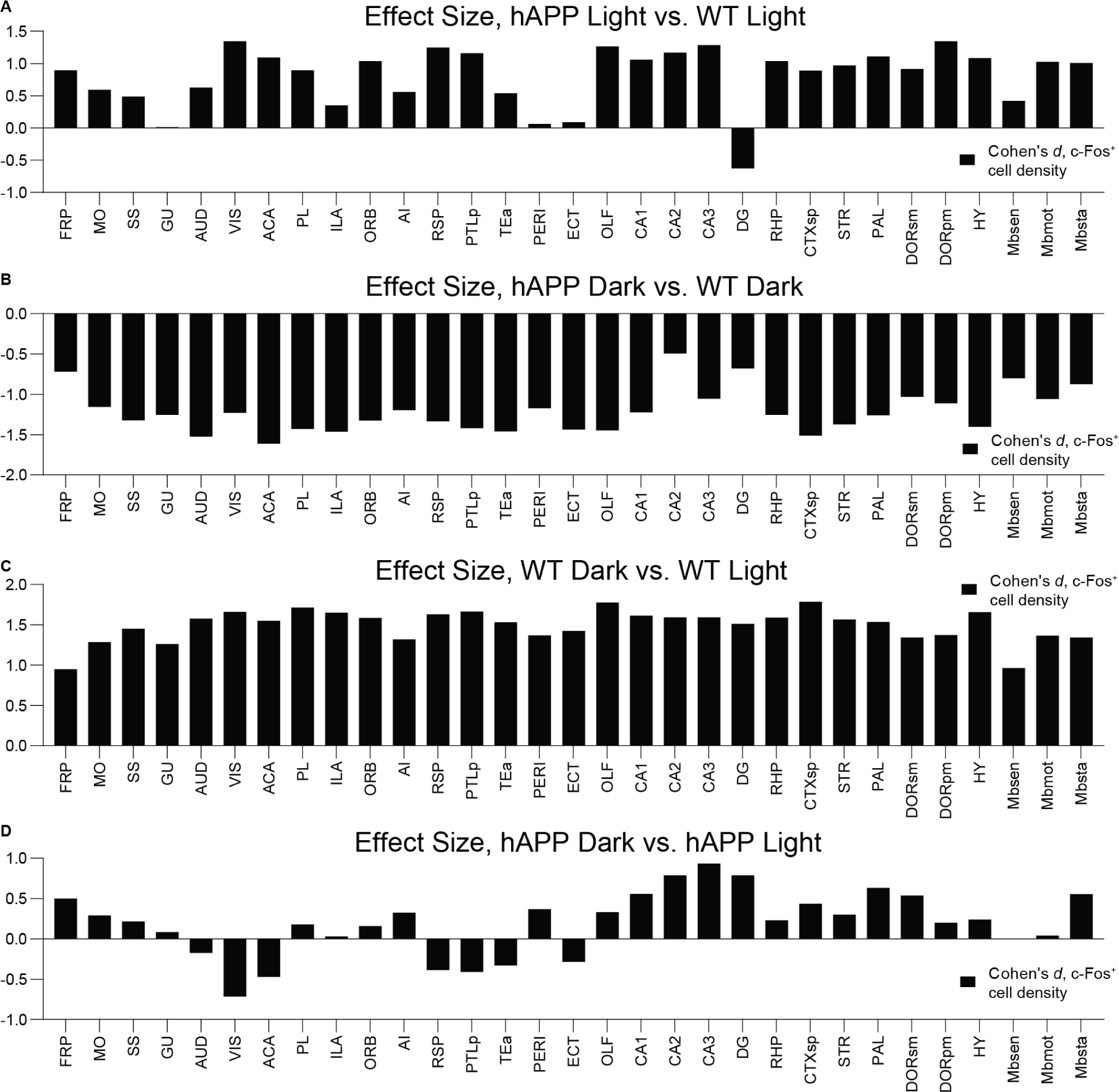
Effect size of hAPP overexpression and light deprivation on c-Fos^+^ cell density across the brain. **A-B)** Effect size (Cohen’s *d*) of hAPP on c-Fos^+^ cell density for mice housed for one week under a normal 12h light/dark cycle (A) or 24h dark (B). **C-D)** Effect size of one week of 24h dark housing on WT (C) and hAPP mice (D). n = 8 WT light, 9 WT dark, 10 hAPP light, and 8 hAPP dark housed mice.

**Supplementary Figure 6:**
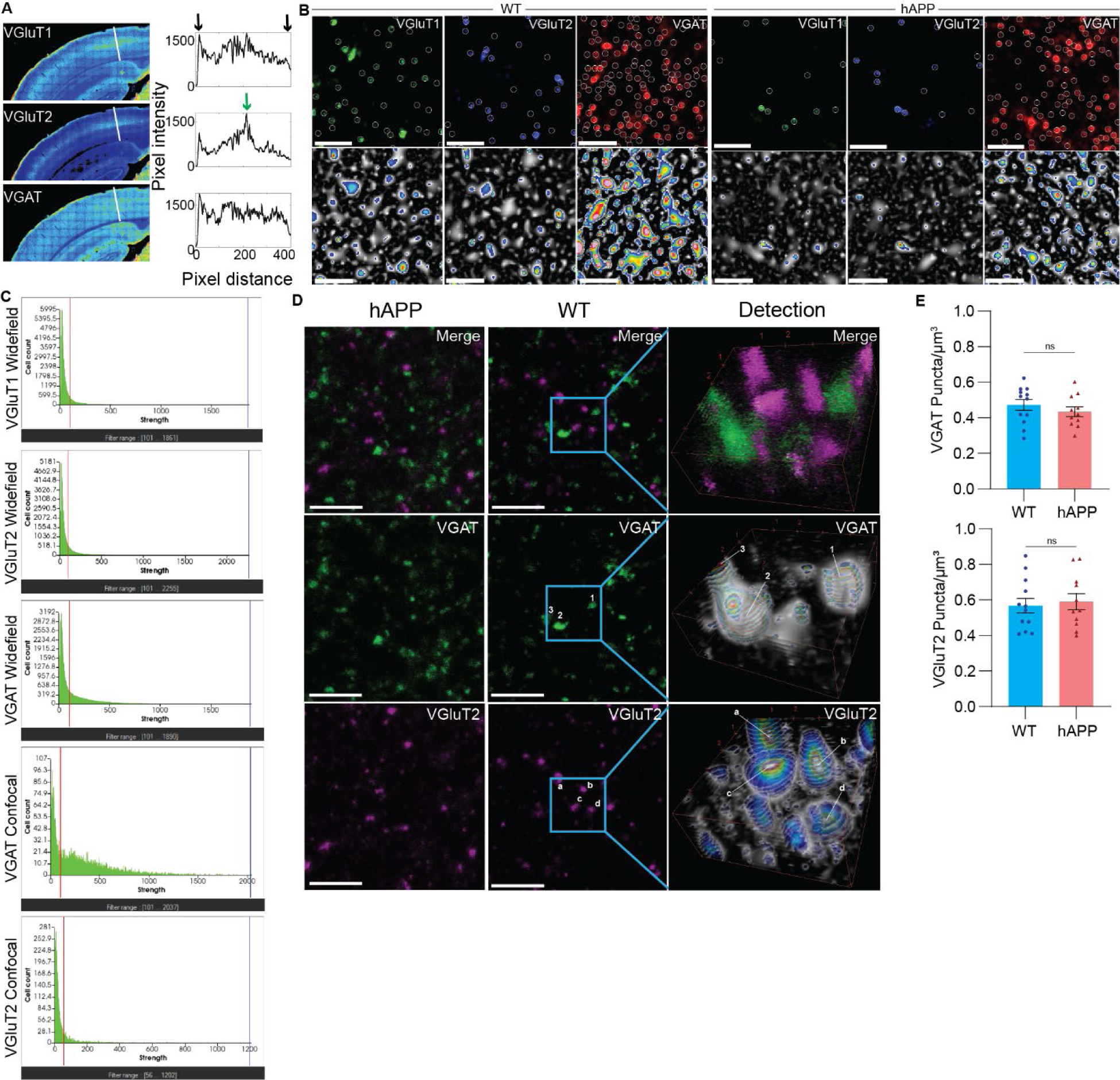
Visualization of synaptic marker expression patterns, synapse detection, and puncta clustering. **A)** Representative pseudocolored images of VGluT1, VGluT2, and VGAT labeling patterns. The warmer color indicates higher pixel intensity. The display range is kept identical for all three images. The white line spans all layers of the visual cortex, and the corresponding pixel intensity is shown on the right. Pixel distance 0 is right outside the beginning of the slice and ends in the fiber tract beneath layer 6 (indicated by black arrows). The green arrow in the VGluT2 image (middle) shows the characteristic increase in VGluT2 puncta from the thalamus to layer 4 of the visual cortex. **B)** Representative images showing synaptic puncta detection output (top panels) and LoG-filtered images of the same field of view in the same channel (bottom panels). For LoG images, pixels under the detection threshold (LoG = 101) are shown in greyscale, and pixels above the detection threshold are shown in a 14-color heatmap. Scale bar = 5 µm. **C)** Representative histograms showing the number of detected objects meeting the size criteria for synaptic puncta (0.32-1.97 µm diameter) at each LoG threshold (Strength) for each cell or synapse detection threshold. Thresholds used for detection (red line) were set based on the typical elbow point in WT animals, below which point many dim objects (false positives) would be detected. The x-axes span the full ranges of the data, though some high LoG value data points are not visible due to the large number of detected “objects” at the low end. **D)** Representative images of VGAT and VGluT2 signal from high-resolution confocal imaging of the primary visual cortex. Merged (upper) and individual channel (center, lower) top-down views through representative 3D reconstructions for hAPP (left) and WT mice (center). The area in the blue boxes for WT reconstructions are shown as merged 5×5×2 µm 3D reconstructions (upper right) and as individual channels with respective volumetric LoG filters applied (center and lower right). For LoG images, pixels under the detection threshold are shown in greyscale, and pixels above the detection threshold are shown in a 14-color heatmap. LoG heatmaps are shown on individual planes. Only 3D objects meeting the LoG and expected synaptic puncta size criteria are included in analysis. Example puncta showing above-threshold LoG values for VGAT (1-3, center right) and VGluT2 (a-d, lower right) channels. Scale bar = 5 µm. **E)** VGAT and VGluT2 puncta density from high-resolution primary visual cortex z-stacks. Data are presented as mean ± SEM. ns, not significant (p > 0.05), student’s *t* tests, n = 12 WT, 11 hAPP mice.

**Supplementary Table 1:** Abbreviations, full structure names, and pooled anatomical brain area for all annotated brain regions from the Allen CCFv3.

| **Abbreviation** | **Full Structure Name** | **Pooled Structure** |
| --- | --- | --- |
| FRP | Frontal pole, cerebral cortex | Isocortex |
| MO | Somatomotor areas | Isocortex |
| SS | Somatosensory areas | Isocortex |
| GU | Gustatory areas | Isocortex |
| AUD | Auditory areas | Isocortex |
| VIS | Visual areas | Isocortex |
| VISal | Anterolateral visual area | Isocortex |
| VISam | Anteromedial visual area | Isocortex |
| VISl | Lateral visual area | Isocortex |
| VISp | Primary visual area | Isocortex |
| VISp2/3 | Primary visual area, layer 2/3 | Isocortex |
| VISp4 | Primary visual area, layer 4 | Isocortex |
| VISp5 | Primary visual area, layer 5 | Isocortex |
| VISp6a | Primary visual area, layer 6a | Isocortex |
| VISp6b | Primary visual area, layer 6b | Isocortex |
| VISpl | Posterolateral visual area | Isocortex |
| VISpm | Posteromedial visual area | Isocortex |
| VISli | Laterointermediate area | Isocortex |
| VISpor | Postrhinal area | Isocortex |
| ACA | Anterior cingulate area | Isocortex |
| PL | Prelimbic area | Isocortex |
| ILA | Infralimbic area | Isocortex |
| ORB | Orbital area | Isocortex |
| AI | Agranular insular area | Isocortex |
| RSP | Retrosplenial area | Isocortex |
| PTLp | Posterior parietal association areas | Isocortex |
| VISa | Anterior area | Isocortex |
| VISrl | Rostrolateral visual area | Isocortex |
| TEa | Temporal association areas | Isocortex |
| PERI | Perihinal area | Isocortex |
| ECT | Ectorhinal area | Isocortex |
| OLF | Olfactory Areas | Olfactory Areas |
| CA1 | Field CA1 | Hippocampal Formation |
| CA2 | Field CA2 | Hippocampal Formation |
| CA3 | Field CA3 | Hippocampal Formation |
| DG | Dentate gyrus | Hippocampal Formation |
| RHP | Retrohippocampal region | Hippocampal Formation |
| CTXsp | Cortical subplate | Cortical Subplate |
| STR | Striatum | Cerebral Nuclei |
| PAL | Pallidum | Cerebral Nuclei |
| dLGN | Dorsal part of the lateral geniculate complex | Interbrain |
| DORsm | Thalamus, sensory-motor cortex related | Interbrain |
| DORpm | Thalamus, polymodal association cortex related | Interbrain |
| LGN | Lateral geniculate complex | Interbrain |
| LP | Lateral posterior nucleus of the thalamus | Interbrain |
| RT | Reticular nucleus of the thalamus | Interbrain |
| HY | Hypothalamus | Hypothalamus |
| Mbsen | Midbrain, sensory related | Midbrain |
| SC | Superior colliculus | Midbrain |
| Mbmot | Midbrain, motor related | Midbrain |
| Mbsta | Midbrain, behavioral state related | Midbrain |

